# Characterizing Molecular and Synaptic Signatures in mouse models of Late-Onset Alzheimer’s Disease Independent of Amyloid and Tau Pathology

**DOI:** 10.1101/2023.12.19.571985

**Authors:** Kevin P. Kotredes, Ravi S. Pandey, Scott Persohn, Kierra Elderidge, Charles P Burton, Ethan W. Miner, Kathryn A. Haynes, Diogo Francisco S. Santos, Sean-Paul Williams, Nicholas Heaton, Cynthia M. Ingraham, Christopher Lloyd, Dylan Garceau, Rita O’Rourke, Sarah Herrick, Claudia Rangel-Barajas, Surendra Maharjan, Nian Wang, Michael Sasner, Bruce T. Lamb, Paul R. Territo, Stacey J. Sukoff Rizzo, Gregory W. Carter, Gareth R. Howell, Adrian L. Oblak

**Author notes:** First authors (Kevin P. Kotredes, Ravi S. Pandey).

## Abstract

**INTRODUCTION:** MODEL-AD is creating and distributing novel mouse models with humanized, clinically relevant genetic risk factors to more accurately mimic LOAD than commonly used transgenic models.

**METHODS:** We created the LOAD2 model by combining APOE4, Trem2*R47H, and humanized amyloid-beta. Mice aged up to 24 months were subjected to either a control diet or a high-fat/high-sugar diet (LOAD2+HFD) from two months of age. We assessed disease-relevant outcomes, including in vivo imaging, biomarkers, multi-omics, neuropathology, and behavior.

**RESULTS:** By 18 months, LOAD2+HFD mice exhibited cortical neuron loss, elevated insoluble brain Aβ42, increased plasma NfL, and altered gene/protein expression related to lipid metabolism and synaptic function. In vivo imaging showed age-dependent reductions in brain region volume and neurovascular uncoupling. LOAD2+HFD mice also displayed deficits in acquiring touchscreen-based cognitive tasks.

**DISCUSSION:** Collectively the comprehensive characterization of LOAD2+HFD mice reveal this model as important for preclinical studies that target features of LOAD independent of amyloid and tau.

## Background

Late-onset Alzheimer’s disease (LOAD2) is the most common form of dementia, caused by a combination of genetic and environmental factors^1-3^. Despite recent approval of anti-amyloid therapies such as Aduhelm®^4^ and Leqembi®^5^, additional therapeutic options are essential to prevent or slow cognitive decline in most cases of LOAD. To achieve this, preclinical models that more faithfully reproduce the complex features of LOAD are required to maximize translatability of preclinical studies to the clinic.

The MODEL-AD (Model Organism Development and Evaluation for Late-Onset Alzheimer’s disease) consortium is charged with creating and phenotyping new mouse models based on the genetics of LOAD^6^. The IU/JAX/PITT MODEL-AD Center has focused on creating models on the C57BL/6J (B6J) genetic background that incorporate the e4 allele of the apolipoprotein E gene (*APOE4*)^7^, the greatest genetic risk factor for LOAD. In addition, we created the e3 (neutral) allele (*APOE3*)^7^. These humanized APOE alleles allow for the unrestricted use and breeding that was not readily available with previous versions^7^. This allowed us to determine the effects of combining multiple genetic risk factors for LOAD. We generated and characterized B6J mice that were double homozygous for both *APOE4* and the *R47H* variant in *Trem2* (Triggering receptor expressed on myeloid cells 2). These mice were termed LOAD1^8^. Although LOAD1 mice did not develop classic hallmarks of LOAD, such as amyloid pathology, neurodegeneration, and cognitive decline, they did show alterations in gene expression levels in the brain similar to those seen in LOAD patients, as well as changes in cerebrovascular blood flow and glucose uptake^8^.

We now present the comprehensive characterization of LOAD2 (B6J.*APOE4*.*Trem2*R47H*.*hA*β triple homozygous), where the Aβ sequence of the mouse *App* gene of LOAD1 mice has been humanized^8^. Data support that the human Aβ sequence is more amyloidogenic than the mouse version and so we test the hypothesis that LOAD2 mice will develop features of LOAD that were absent in LOAD1 mice. In addition, we evaluate a high-fat diet/high-sugar diet (HFD), a common environmental stressor, that human and mouse studies show increases risk for LOAD^9, 10^. For instance, our previous study showed chronic consumption of a HFD exacerbated the genetic effects of LOAD1 mice carrying the *Plcg2*M28L^10^*. Here, using a combination of a cross-sectional and longitudinal design, cohorts of male and female LOAD1 and LOAD2 mice were fed either a control diet (CD) or HFD from 2 months of age and evaluated at 4, 12, 18 or 24 months of age. Data show that unlike LOAD1 mice, LOAD2 mice fed a HFD (LOAD2+HFD) resulted in age-related neurodegeneration, cognitive deficits, elevations in insoluble Aβ and LOAD-relevant imaging abnormalities, and increased neurofilament light chain (NfL) in the plasma. We propose LOAD2+HFD as a relevant mouse model for investigating therapeutic interventions independent of targeting tau and amyloid pathologies.

## Methods

### *Creation of the humanized A*β allele

The humanized Aβ allele was created by direct delivery of CRISPR-Cas9 reagents to mouse zygotes of the APOE4/Trem2*R47H model, (B6(SJL)-*Apoe^tm1.1(APOE*4)Adiuj^ Trem2^em1Adiuj^*/J or “LOAD1”, JAX #28709, https://www.jax.org/strain/028709) which was previously described^8^

Analysis of genomic DNA sequence surrounding the target region, using the Benchling (www.benchling.com) guide RNA design tool, identified a gRNA sequence (TTTGATGGCGGACTTCAAATC) with a suitable target endonuclease site in exon 14 of the mouse App locus. Streptococcus pyogenes Cas9 (SpCas9) V3 protein and gRNA were purchased as part of the Alt-R CRISPR-Cas9 system using the crRNA:tracrRNA duplex format as the gRNA species (IDT, USA). Alt-R CRISPR-Cas9 crRNAs (Product# 1072532, IDT, USA) were synthesized using the gRNA sequences specified in the DESIGN section and hybridized with the Alt-R tracrRNA (Product# 1072534, IDT, USA) as per manufacturer’s instructions. A single-stranded DNA repair construct (synthesized by Genscript) with the sequence 5’- CTGGGCTGACAAACATCAAGACGGAAGAGATCTCGGAAGTGAAGATGGATGCAGA ATTC**C**GACATGATTCAGGAT**A**TGAAGTCC**AT**CATCAAAAACTGGTAGGCAAAAATAAACTGCCTCTCCCCGAGATTGCGTCTGGCCAGATGAAAT-3’ was used to introduce the G601R, F606Y, and R609H amino acid changes in the mouse *App* sequence (corresponding to G676R, F681Y and R864H in human *APP*) such that the Ab-42 region matches the human sequence (Figure 1A).

**FIGURE 1:**
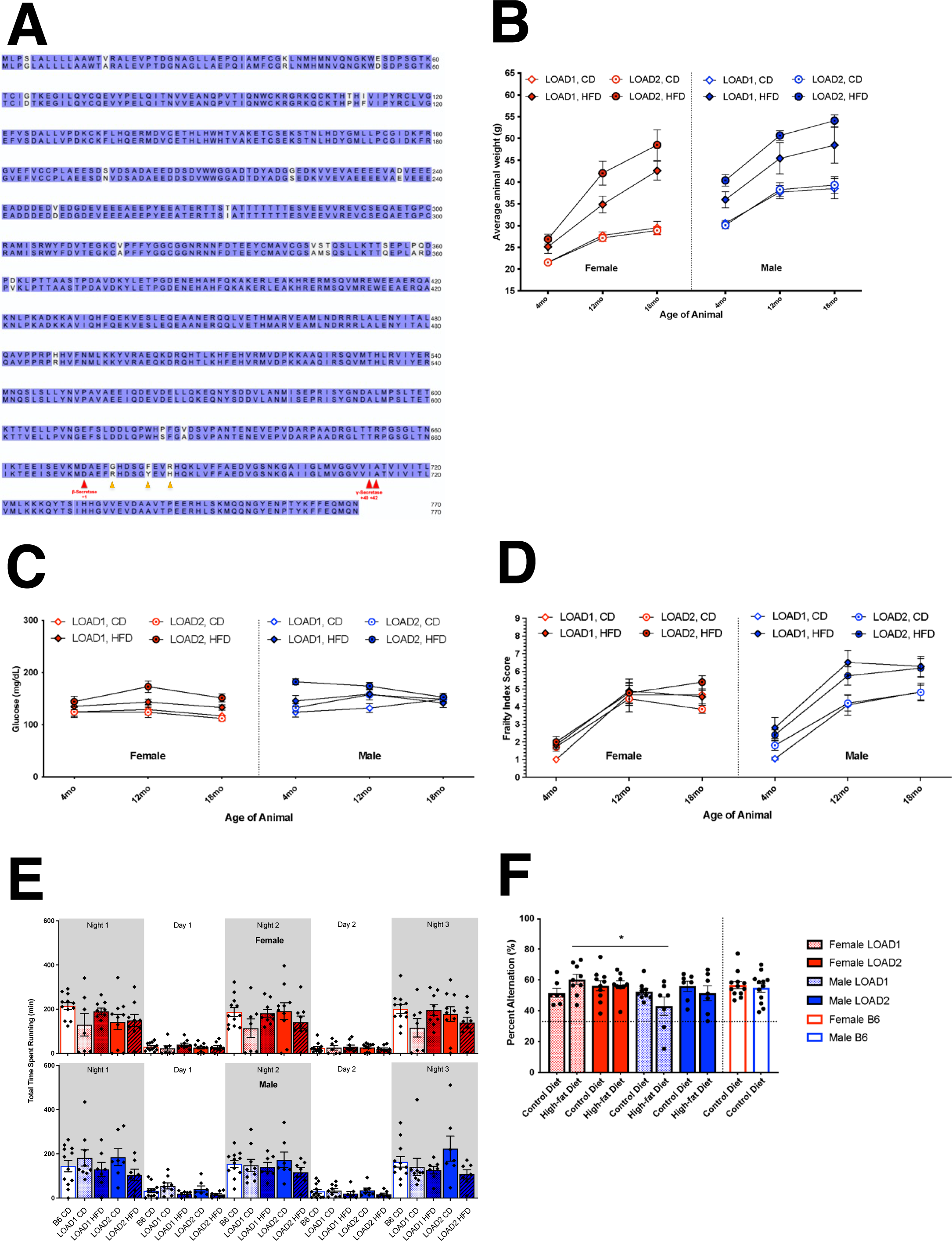
Longitudinal metabolic and behavioral phenotyping of mice on high-fat diet. LOAD1 (*APOE4/Trem2*R47H*) and LOAD2 (*hAbeta/APOE4/Trem2*R47H*) animal strains differ in the *App* allele with a humanized Abeta1-42 region (G601R, F606Y, R609H in the mouse gene, corresponding to amino acid positions 676, 681, 684 in the human *APP* locus) (A). Alignment of mouse (top; Uniprot ID P12023) and humanized (bottom; Uniprot ID P05067) amyloid precursor protein (APP) amino acid sequences. White letters denote non-homology. Red arrows indicate cleavage sites of processing enzymes. Yellow arrows denote sites of humanizing mutations in *App* allele (LOAD1, top, and LOAD2, bottom). (Cohort 1) Animals of an 18-month longitudinal cohort were assayed at 4-, 12-, and 18-months of age. Males and females, of LOAD1 and LOAD2 genotypes, fed either control diet (CD) or high-fat diet (HFD) beginning at 2-months of age were measured for body weight (B), fasted blood glucose (C), and frailty assay index score (D), as a measure of general animal health changes. Running wheel assay measured average animal activity time for three days and nights at the 18-month age timepoint (E). Spontaneous alternation behavioral assay was utilized to measure cognition longitudinally across ages at the 18-month age timepoint (F). (Three-way ANOVA [sex, genotype, diet effects]; *=p<0.05)

Founders were bred to the LOAD1 model and genotyped for the humanized Aβ locus by PCR using forward primer 5’-CAGTTTTTGCCTCCTTGTGG-3’ and reverse primer 5’-GGCTTCTGCTCAGCAAGAACTA-3’. A positive reaction was determined by the presence of a band of 362bp. The resulting strain (“LOAD2”) is available as JAX #30670, B6J.Cg- *Apoe^tm1.1(APOE*4)Adiuj^ App^em1Adiuj^ Trem2^em1Adiuj^*/J (https://www.jax.org/strain/030670; B6J.*APOE^E4/E4^*.*Trem2^R47H/R47H^*.*App*^hAβ/hAβ^). The *App* allele alone is available as JAX #33013, B6J.Cg- *App^em1Adiu^*^j^/J (https://www.jax.org/strain/033013).

### Cohort generation and evaluation

To evaluate LOAD-relevant phenotypes, five cohorts of LOAD2 mice and controls were created at The Jackson Laboratory (JAX, cohort 1), Indiana University (IU, cohorts 2 and 3) and University of Pittsburgh (PITT, cohorts 4 and 5). Breeding, mouse husbandry, and assays common across sites were standardized as much as possible. Below, we provide brief details of each cohort and assays performed with full details included provided in Supplemental Methods.

### Cohort 1 – The Jackson Laboratory

All procedures were approved by The Jackson Laboratory Institutional Animal Care and Use Committee (IACUC).

#### Experimental groups

To create experimental groups, B6J.*APOE^E4/E4^*.*Trem2^R47H/R47H^*.*App*^hAβ/+^ mice were intercrossed to create B6J.*APOE^E4/E4^*.*Trem2^R47H/R47H^*.*App*^hAβ/hAβ^ (LOAD2) and B6J.*APOE^E4/E4^*.*Trem2^R47H/R47H^*.*App*^+/+^ (LOAD1) control mice. In appreciation of sexual dimorphism observed in human aging and disease, four groups of male and female mice were established for a combination of longitudinal and cross-sectional phenotyping at 4-, 12-, 18-, and 24-months. The 18-months group was assessed for biometrics and plasma biomarkers at 4-, 8-, 12- and 18-months.

All mice were initially provided LabDiet® 5K52/5K67 (6% fat; control diet, CD). At 2 months of age, each experimental group was randomized into two groups, the control group, and the high-fat diet (HFD) group. The control groups continued on CD *ad libitum*, while the HFD groups were provided ResearchDiet^®^ feed D12451i (45% high fat, 35% carbohydrates) *ad libitum*. Due to attrition between 18-24 months of age, the 24-month HFD cohort was not sufficiently powered and so not analyzed. For *in vivo* studies – at least 10 mice/sex/genotype/age/diet were evaluated. For post-mortem analyses, 6 mice/sex/genotype/age/diet were evaluated unless otherwise stated.

#### Phenotyping

The cross-sectional phenotyping battery included *in vivo* frailty, behavioral phenotyping, and metabolic profiling and biomarker (e.g., Neurofilament light chain, NfL) analyses in the plasma. Postmortem brain tissue was examined for transcriptomic and proteomic analyses as well as neuropathological indications of disease (amyloid, neuronal cell loss, and glial activation). For full details see Supplemental Methods (Supplemental Figure 1).

### Cohorts 2: Indiana University

All procedures were approved by the Indiana University Institutional Animal Care and Use Committee (IACUC). The same bedding, light cycle, and water conditions as The Jackson Laboratory were used at Indiana University.

#### Experimental groups

LOAD2 mice were imported from JAX and bred at IU. LOAD2 mice were initially crossed to LOAD1 mice to create B6J.*APOE^E4/E4^*.*Trem2^R47H/R47H^*.*App*^hAβ/+^ mice that were then intercrossed to create B6J.*APOE^E4/E4^*.*Trem2^R47H/R47H^*.*App*^hAβ/hAβ^ (LOAD2) mice. One group of at least 10 male and 10 female mice were established for longitudinal phenotyping at 4, 12, and 18 months. Similar to the JAX cohort, mice were initially provided CD before half the mice in each group were switched to HFD.

#### Phenotyping

At 4, 12, and 18 months, mice underwent *in vivo* MR imaging (T2 weighted images) and blood draws for biomarker (e.g., Aβ species, cytokines) analyses. For full details see Supplemental Methods. At 18 months, tissues were collected as described for Cohort 1.

### Cohort 3: Indiana University

All procedures were approved by the Indiana University Institutional Animal Care and Use Committee (IACUC). The same bedding, light cycle, and water conditions as The Jackson Laboratory were used at Indiana University.

#### Experimental groups

Experimental groups of male and female LOAD2 mice on HFD or CD were established as described for Cohort 3 (n=12 mice/sex/genotype/age/diet). Three groups were established for cross-sectional analyses at 4, 12, and 18 months.

#### Phenotyping

To evaluate neurovascular uncoupling, *in vivo* PET/CT imaging was performed on all mice measuring regional blood flow (via ^64^Cu-pyruvaldehyde-bis(N4- methylthiosemicarbazone, ^64^Cu-PTSM) and regional glycolytic metabolism (via 2-^18^F-2- deoxyglucose, ^18^F-FDG). Findings from *in vivo* PET/CT were confirmed using autoradiography. For full details see Supplemental Methods.

### Cohort 4 and 5: University of Pittsburgh

All procedures were approved by the University of Pittsburgh Institutional Animal Care and Use Committee (IACUC). Detailed mouse husbandry, diet restriction, and phenotyping methods are included in the Supplemental Methods.

#### Experimental groups

Two experimental cohorts were evaluated for plasma biomarkers and cognitive testing. Breeding pairs of LOAD2 mice were imported from JAX and bred at the University of Pittsburgh to create Cohort 4. One group of n=18 male and n=18 female mice were established for longitudinal blood plasma collection followed by cognitive assessments using the touchscreen. Subjects were reared on normal control diet (CD) (LabDiet® 5P76) provided *ad libitum* until 2 months of age at which time n=13 male and n=14 female mice were randomly assigned to receive ad libitum HFD (LOAD2+HFD). For cohort 5, LOAD2 mice (B6J.*APOE^E4/E4^*.*Trem2^R47H/R47H^*.*App*^hAβ/hAβ^) were bred with C57BL/6J to provide littermate controls. The F1 offspring which were triple heterozygotes (B6J.*APOE^E4/+^*.*Trem2^R47H/+^*.*App*^hAβ/+^) were then crossbred to produce F2 offspring including the LOAD2 triple homozygote mice (n=9/sex) and triple wildtype littermate controls (n=6/sex). All mice were initially reared on CD, with n=3/sex wildtype controls and n=6/sex LOAD2 switched to ad libitum HFD at 6-12 months of age. Prior to touchscreen testing, mice were individually housed and restricted to 80-85% of free-feeding body weight. Mice were weighed daily and provided a ration of the respective CD or HFD diets that maintained them at 80-85% restriction.

#### Phenotyping

Blood plasma was evaluated longitudinally prior to the start of HFD, followed monthly for analysis of cytokines and every 3 months for analysis of Aβ species. To evaluate the effect of food restriction on plasma biomarker levels, brief 2-week periods of food restriction as described above were administered to Cohort 4 at 8-8.5 months of age and at 10-10.5 months of age. At 14 months of age, Cohort 4 was enrolled in Touchscreen cognitive testing and maintained continuously on dietary restriction until the conclusion of touchscreen testing (Figure 8A); while Cohort 5 began food restriction and touchscreen testing at 11-17 months of age (Supplemental Figure 7A). It is important to note that the present studies used a 10% sucrose solution for the reward which is a departure from the standard touchscreen protocols that use strawberry flavored milkshake-based rewards^11^. We intentionally chose to avoid milk-based rewards given that the constituents of milk and dairy products may contribute to attenuation of AD related pathologies including amyloid deposition and inflammation^12-14^. Notably 10% sucrose is a common and well-established reinforcer for mice in operant based tasks and therefore was a salient alternative as evidenced by the ability of all subjects to demonstrate consumption of the reward and acquire the touch-reward association during the initial phase of the task.

## Results

LOAD2 mice were created at JAX and distributed to IU and PITT for evaluation of LOAD relevant phenotypes using the IU/JAX/PITT MODEL-AD center pipeline that includes a combination of human-relevant *in vivo* and post-mortem assays. The primary goal of the phenotyping pipeline is to determine the utility of new LOAD models for preclinical testing. Five cohorts of LOAD2 and control mice fed either a HFD (high-fat diet) or CD (control diet) were evaluated at JAX (Cohort 1: biometrics, behavior, plasma biomarkers, neuropathology, transcriptomics, proteomics), IU (Cohort 2: MRI, plasma biomarkers and cytokines, biochemistry, neuropathology; Cohort 3: PET/CT, autoradiography), and PITT (Cohorts 4 and 5: longitudinal plasma biomarkers, Touchscreen cognitive testing) (Supplemental Figure 1). Unless otherwise stated, to evaluate LOAD2 phenotypes (on CD or HFD), LOAD1 mice were used as the control genotype to evaluate the effects of humanizing the Aβ sequence in the context of the *APOE4* and Trem2*R47H risk alleles (LOAD1). LOAD1 and LOAD2 mice both express humanized *APOE4* and *Trem2*R47H risk* alleles on a B6J background, however LOAD2 animals also express a humanized allele for *App* (Figure 1A).

### LOAD2 mice show diet- and age-dependent neuronal cell loss and plasma NfL increases (cohort 1)

Cohort 1 comprised four age groups (4, 12, 18 and 24 months). The 4-, 12-, and 18- month groups included LOAD1 and LOAD2 mice on both HFD and CD. The 24-month group included only mice on CD. We first evaluated the 18-month group using *in vivo* assays following a longitudinal design. Longitudinal testing and sampling were performed at 4-, 12-, and 18-months of age. We observed significant, diet-driven increases in body weight with age (Figure 1B). All mice showed an increase in weight with age, but mice fed HFD showed pronounced weight gain until 18 months of age. Females on the HFD displayed significant weight gain from 4- to 12-months, but only made modest gains from 12- to 18- months. Males fed a HFD were more accelerated than females, and LOAD2 males were consistently heavier than LOAD1 males at all timepoints. However, fasted blood glucose measurements did not appear to be age-, diet-, or genotype-dependent, though slightly elevated levels were observed in LOAD2 mice fed HFD (LOAD2+HFD) (Figure 1C). As expected, age was a strong factor of increased frailty^15^ (Figure 1D). However, we did not see significant diet-related changes in frailty scores in females until 18 months of age and only in LOAD2 genotype animals. Males consistently displayed increased frailty driven by diet, but not genotype, as early as 8 months of age.

HFD reduced performance in open field assays compared to CD. Specifically total distance traveled decreased significantly only in LOAD2 animals (Supplemental Figure 2A), but performance was not affected by sex or genotype alone. Differences in total vertical activity were only observed in males, of both genotypes, to be decreased by HFD (Supplemental Figure 2B). Rotarod performance was decreased in a HFD-dependent manner only (Supplemental Figure 2C). LOAD2+HFD animals, however, particularly males, demonstrated a reduction in running wheel activity during the active period (dark cycle) compared to LOAD1 mice or those fed CD (Figure 1E). Hippocampal working memory in the spontaneous alternation assay as a measure of cognitive function was intact across all groups with all subjects performing >chance levels which is calculated as 22% in this assay (Figure 1F).

Brains from mice from the 18-month group were harvested (along with the 4-, 12-, and 24-month groups). One hemisphere (right) from each brain was prepared for transcriptomics and proteomics, while the other hemisphere (left) was evaluated for neuronal cell loss, microglia number, astrocyte reactivity and amyloid plaques in the cortex and hippocampus by immunofluorescence (Supplemental Figure 3). At 18 months of age, neuron counts in the cortex revealed a subtle but statistically significant decrease in NeuN+DAPI+ cells in female LOAD2+HFD compared to female LOAD2+CD and female LOAD1+HFD (Figure 2 A,B). No differences were observed in cortical IBA1+DAPI+ microglia number (Figure 2 C,D) across all groups, and a small but significant decrease in hippocampal astrocyte reactivity (assessed using GFAP+DAPI+) in male LOAD2+HFD mice compared to those fed a CD diet (Figure 2 E,F). ThioS staining revealed no evidence of amyloid plaques in any groups (Figure 2G). To determine whether neuronal cell changes in LOAD2+HFD was reflected by changes in the plasma, NFL, a clinically relevant biomarker^16^ was assessed. Plasma NfL levels were significantly increased in LOAD2 animals on both diets (Figure 2H). Similarly, we saw significant increase in LOAD1+HFD relative to LOAD1+CD.

**FIGURE 2:**
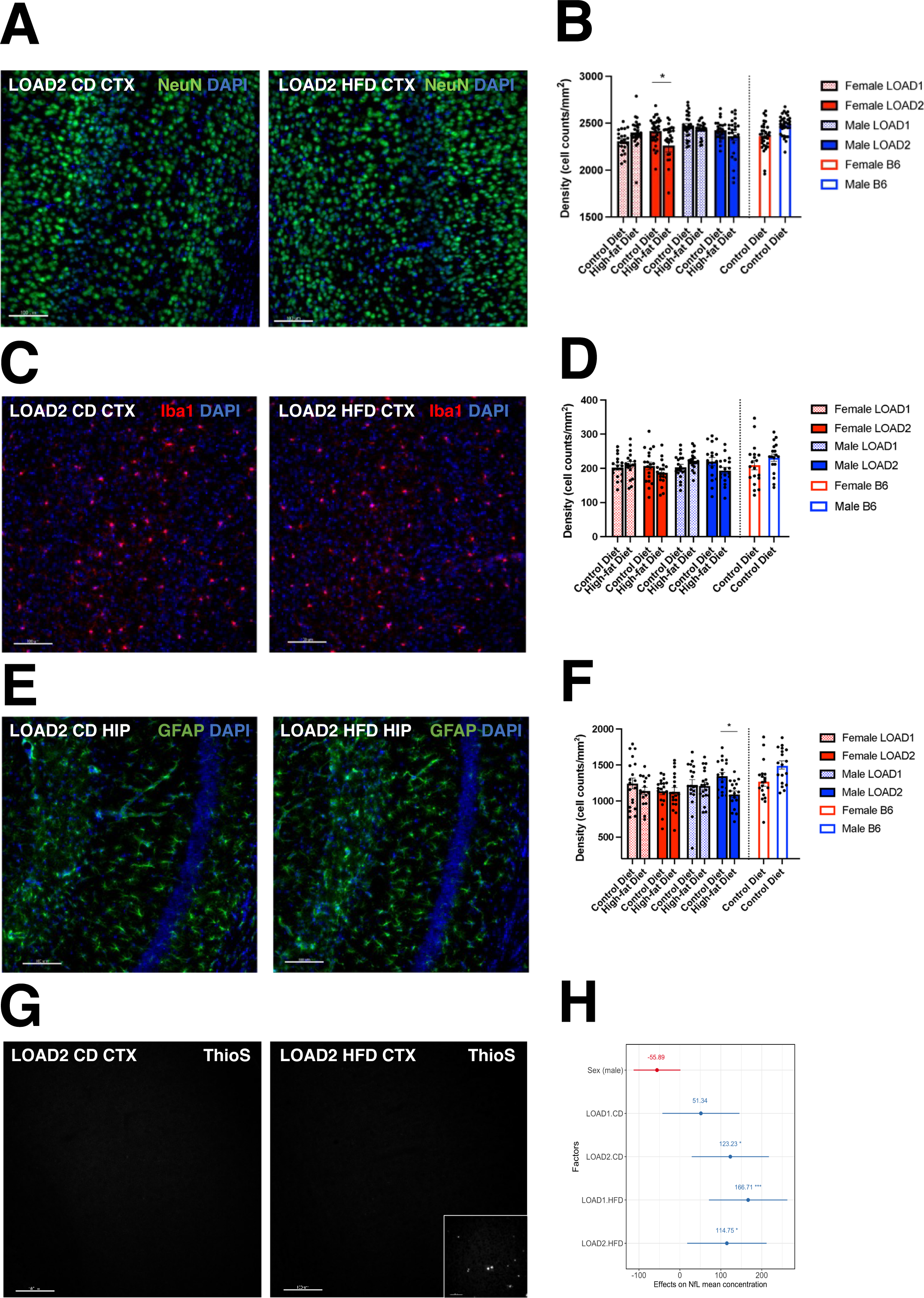
Neuropathological assessment of brain tissue. Immunohistochemistry of brain tissue in the cortex and hippocampus from 18-month-old animals stained for cell markers to reveal genotype- and diet-driven differences in glial cell densities. Slices of brain hemispheres were stained with (A,B) NeuN (neurons) or (C,D) IBA1 (microglia) (representative cortical images shown from LOAD2 females with DAPI co-stain) and counted relative to area. Astrocytes (GFAP) quantitated in the hippocampus of LOAD2 females fed either CD or HFD (E-F). ThioS staining of brain tissue to visualize amyloid plaques (representative images shown from LOAD2 females) (G). Inlay: scaled image of 12mo B6J.*APP-SAA* hyper-amyloid positive controls. LISA testing for neurofilament light-chain (NfL) in plasma derived from terminal, peripheral blood samples at 18-months of age. Linear regression analyses were performed to identify effect of each factor on NfL levels (H). (NeuN=neuronal marker; ThioS=amyloid plaques; GFAP=astrocyte marker; IBA1=microglial marker. Scale bar equals 100μm.)

### Differential analysis identifies strong transcriptional changes in female mice expressing humanized Aβ on high-fat high sugar diet (cohort 1)

To identify molecular effects of humanizing the Aβ peptide, we first performed pairwise differential analysis between LOAD2 and LOAD1 mice at all ages for both sexes. Differential expression analyses identified very few significantly differentially expressed genes (DEGs) (p < 0.05) at 4 and 18 months old LOAD2 mice compared to age and sex-matched LOAD1 mice (Supplementary Table A). At 12 months, there were 57 DEGs (38 upregulated, 19 downregulated; p < 0.05) in male LOAD2 mice, and 17 DEGs (3 upregulated, 14 downregulated; p < 0.05) in female LOAD2 mice (Supplementary Table A). KEGG functional enrichment analysis identified enrichment of “protein processing in ER” in upregulated DEGs in 12 months old LOAD2 male mice and “MAPK signaling pathway” in downregulated DEGs in 12 months old LOAD2 female mice (Supplementary Table B). At 24 months, there were only 5 significantly differentially upregulated genes (p < 0.05) in male LOAD2 mice, and 30 DEGs (12 upregulated, 18 downregulated; p < 0.05) in female LOAD2 mice (Supplementary Table A). Upregulated DEGs in male and female LOAD2 mice were enriched for the “motor proteins” KEGG pathways (Supplementary Table B).

Next, we performed differential analyses in 18-month-old LOAD2 and LOAD1 mice compared to age and sex-matched B6J control mice. In females, we observed only 9 significantly DEGs (3 upregulated, 6 downregulated) (p < 0.05) in LOAD2 mice on control diet (CD), while 2988 genes were significantly differentially expressed (1565 upregulated, 1423 downregulated) (p < 0.05) in LOAD2+HFD (Supplementary Table A). We observed 44 significantly DEGs (19 upregulated, 25 downregulated) (p < 0.05) in female LOAD1+CD, while 164 genes were significantly differentially expressed (117 upregulated, 47 downregulated) (p < 0.05) in female LOAD1+HFD (Supplementary Table A). In males, we observed a total of 7 and 23 significantly DEGs (p < 0.05) in LOAD1 and LOAD2 mice on CD, respectively, while 39 and 98 genes were significantly expressed (p < 0.05) in LOAD1 and LOAD2 mice on HFD, respectively (Supplementary Table A). Overall, we observed more differentially expressed genes in mice conditioned on HFD and this effect was more prominent in female mice expressing humanized Aβ.

Functional enrichment analyses of DEGs in female LOAD2+HFD identified enrichment of multiple KEGG pathways such as “glutamatergic synapse”, “dopaminergic synapse”, and “MAPK signaling pathway” in upregulated genes, while downregulated genes were enriched for KEGG pathways such as “lysosome”, “fatty acid metabolism”, “TCS cycle”, and “valine, leucine and isoleucine degradation”. Differentially upregulated genes in female LOAD1+HFD were enriched for “circadian entrainment”, while downregulated genes were enriched for “phagosome” KEGG pathway (Supplementary Table B). We did not observe enrichment for any KEGG pathways in DEGs in LOAD1 and LOAD2 mice on control diet.

Next, we assess the effect of HFD by performing differential analysis between mice fed the HFD with age, sex, and genotype-matched mice on CD. We observed 260 significantly DEGs (154 upregulated, 106 downregulated) (p < 0.05) in female LOAD2+HFD compared to female LOAD2+CD, while 45 significantly DEGs (9 upregulated, 36 downregulated) (p < 0.05) in male LOAD2+HFD compared to male LOAD2+CD (Supplementary Table A). In LOAD1 male mice, we observed a total of 12 DEGs (p < 0.05) on HFD compared to CD, while only 2 DEGs (p < 0.05) on HFD compared to CD in female LOAD1 mice (Supplementary Table A). Upregulated genes in female LOAD2 female mice on HFD compared to CD were enriched for KEGG pathways such as “glutamatergic synapse”, “dopaminergic synapse”, and “MAPK signaling pathway”, while downregulated genes in female LOAD2+HFD compared to CD were enriched for “motor proteins” pathway (Supplementary Table B).

### Gene modules associated with AD pathology driven by age and high-fat high-sugar diet

Differential expression analyses identified subtle changes at the gene level and suggested pronounced effect of LOAD2 genotype by high-fat/high-sugar diet (HFD) in aged mice. To further ensure these signals, we performed a weighted gene co-expression network analysis (WGCNA) ^17^ on the brain transcriptome to identify gene expression changes in a system-level framework. WGCNA identified 30 distinct modules of co-expressed genes (Supplementary Table B2).

To understand the functional significance of these modules, we correlated each module eigengene with age, sex, diet, genotype, and measured behavioral assays such as cumulative frailty score, body weights, NfL, plasma cytokines, GFAP, Iba1 and NeuN counts (Supplemental Figure 4). Eighteen of these 30 modules were significantly correlated with age (p < 0.05) (turquoise, lightyellow, darkgreen, green, darkgrey, floralwhite, purple, brown4, red, skyblue3, lightcyan, orangred4, darkorange2, plum1, orange, lightcyan1, greenyellow, and blue). Six modules were significantly correlated with HFD (p < 0.05) (lightyellow, turquoise, brown4, darkgrey, darkred, orangered4). Seven modules were significantly correlated with sex (p < 0.05) (sienna3, greenyellow, orangered4, darkmagenta, skyblue3, red, bisque4). Two modules (greenyellow and sienna3) were significantly correlated with the Apoe4.Trem2*R47H allele combination in both LOAD1 and LOAD2 (p < 0.05), while five modules were significantly correlated with humanized Aβ specific to LOAD2 (p < 0.05) (brown, thistle3, greenyellow, blue, and orange) (Supplemental Figure 4).

We observed that HFD was strongly significantly correlated with the lightyellow module (r=0.45; p = 9 x 10^−9^), while age was strongly significantly correlated with both the turquoise (r=0.81; p = 2 x 10^−35^) and lightyellow modules (r=0.52; p = 1 x 10^−11^) (Supplemental Figure 4). To further elucidate the association of the lightyellow and turquoise modules with age, sex, and genotype-diet combinations, we performed linear regression analysis using module eigengene as the dependent variable. We determined that the lightyellow module was significantly positively correlated with LOAD1 and LOAD2 genotype with HFD (p < 0.05) and age (p < 0.001), while turquoise module was significantly positively correlated with age (p < 0.001) (Figure 3A). In summary, we observed age and genotype-by-diet effects on the lightyellow module, while the turquoise module is driven only by age. We also observed that the diet and age driven lightyellow module was significantly correlated (p < 0.05) with multiple assays such as NfL, frailty score, plasma cytokines (IL1β, IL10, IL5, IL6, KC-GRO) (Figure 3B, Supplemental Figure 4). In contrast, the age driven turquoise module was significantly correlated (p < 0.05) with effect of age on behavior and weakly correlated with a few plasma cytokines (IL2, IFN-γ) and inflammatory cell counts (Iba1 and GFAP counts) (Figure 3B, Supplemental Figure 4). The lightyellow module was uniquely driven by both age and diet and demonstrated strong positive associations with AD biomarkers such as NfL and multiple cytokines.

**FIGURE 3:**
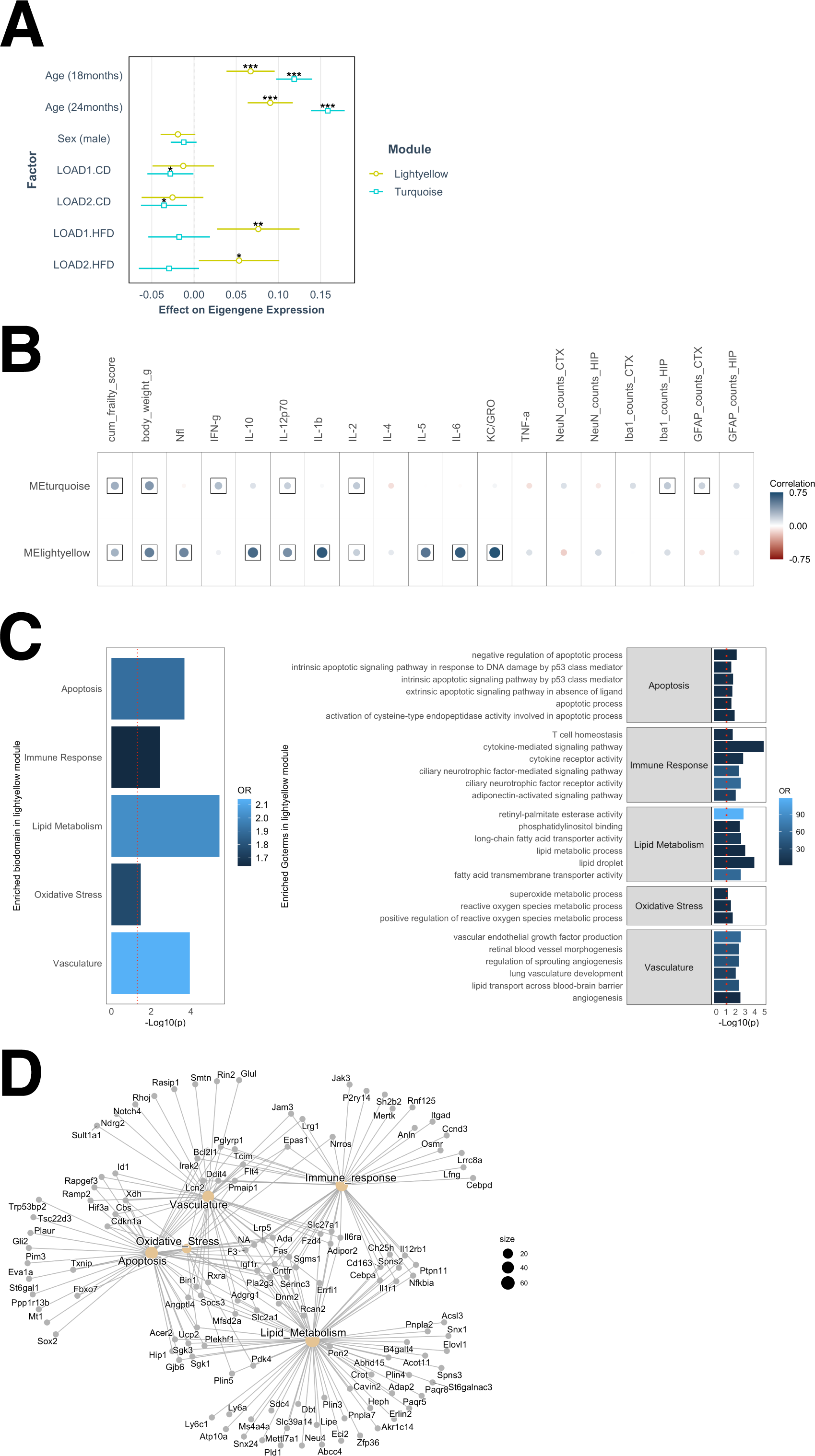
A gene module associated with AD biomarkers is driven by age and high-fat high-sugar diet. The lightyellow gene module was associated with advanced ages (p < 0.001) and both genotype on HFD (p < 0.05), while the turquoise module was associated with age only (p < 0.001) (A). Correlations between the turquoise and lightyellow module eigengenes. Lightyellow was significantly correlated with frailty score, body weight, NfL, and many plasma cytokines (IL-1β, IL-2, IL-12p70, IL-10, IL-5, IL-6, KC-GRO), while the age-driven turquoise module was correlated with behavioral assay (frailty score and body weights) and weakly correlated with a few plasma cytokines (IL-2, IFNγ) and inflammatory cell counts (IBA1 and GFAP counts) (B). Positive correlation coefficients are shown in blue and negative correlations in red, proportional to color intensity and circle size, with frames for significant correlations (FDR < 0.05). AD-related biological domain enrichment analysis in the age and HFD driven lightyellow module gene set using Fisher exact test, with the top six enriched GO terms within each enriched bidomain (C). Network of genes in each enriched biological domain and the lightyellow module (D).

We further assessed the enrichment of AD biological domains^18^ in gene modules. We found that genes in lightyellow modules were significant enriched for the Apoptosis (odds ratio=1.90, p = 2.11 x 10^−4^), Immune Response (odds ratio=1.63, p = 3.67 x 10^−3^), Lipid Metabolism (odds ratio=2.01, p = 3.64 x 10^−6^), Oxidative Stress (odds ratio=1.76, p = 3.32 x 10^−2^), and Vasculature (odds ratio=2.14, p = 1.14 x 10^−4^) AD biological domains (Figure 3C-D; Supplementary Table B3). We also identified GO-terms associated with these biological domains that are significantly enriched in lightyellow gene modules (Figure 3C-D, Supplementary Table B3). On the other hand, genes in the turquoise module were prominently enriched for the Immune Response biological domain (odds ratio=2.39, p = 1.8 x 10^−49^) (Supplementary Table B3). These data suggest that age is strongest risk factor for driving inflammatory changes, while diet effects in aged LOAD mice are associated with multiple AD endophenotypes such as lipid metabolism, immune response, and oxidative stress.

### LOAD mice display proteomics changes characteristic of AD

Tandem mass tag proteomics was performed on hemibrains from LOAD1 and LOAD2 mice at 4, 12, and 18 months of age on control diet. LOAD1 and LOAD2 mice fed HFD were also assayed at 18 months, paired with B6J controls on CD. A total of 10,406 proteins were quantified across 106 samples. To focus on effects in aged mice, we assessed protein expression changes in at 18-month-old LOAD1 and LOAD2 mice on CD and HFD compared to age-matched B6J mice on CD by one-way ANOVA with post-hoc Tukey significance testing (Supplementary Table C). In LOAD1 mice on CD, we observed 1666 significantly differentially expressed proteins (838 upregulated, 828 downregulated) (padj < 0.05), while a total of 2590 proteins were significantly differentially expressed (1237 upregulated, 1353 downregulated) (padj < 0.05) in LOAD1+HFD compared to B6J mice on CD (Supplementary Table C). In LOAD2+CD, we observed a total of 1102 significantly differentially expressed proteins (535 upregulated, 567 downregulated) (padj < 0.05) while in LOAD2+HFD we observed 1839 significantly differentially expressed proteins (897 upregulated, 942 downregulated) (padj < 0.05). We therefore observed hundreds of differentially expressed proteins in both LOAD1 and LOAD2, with greater numbers for both strains on HFD.

To assess disease relevant aspects of mouse models, we performed a correlation analysis between mouse models and 44 human proteomics modules from a LOAD study of the dorsolateral prefrontal cortex^19^. These modules were functionally annotated and named based on protein enrichments and each was assessed for eigenprotein correlations to AD traits including neuropathological markers and cognitive outcomes^19^. Twelve modules were significantly correlated to one or more traits, referred to here as AD modules^19^. We compared protein expression changes in LOAD1 and LOAD2 mice relative to B6J mice at 18 months with changes observed in human AD subjects versus controls for each human protein module. This procedure allowed module-wide assessment of coordinated protein changes and therefore determined murine reproduction of each module that characterizes human LOAD.

LOAD1 and LOAD2 mice were significantly and positively correlated (p < 0.05) with multiple common human AD modules. These included M1_Synapse_Neuron, M3_Oligo_Myelination, and M12_Cytoskeleton (Figure 4A; Supplementary Table D). The M2_Mitochondria module exhibited significant positive correlation (p < 0.05) with all mouse models except LOAD2 mice on HFD, for which a positive correlation did not reach significance (p = 0.06) (Figure 4A; Supplementary Table D). Additional correlations reaching significance included LOAD1 mice on HFD and LOAD2 mice on both diets with M22_Post-Synaptic_Density and M38_Heat_Shock_Folding modules (Figure 4A; Supplementary Table D). LOAD2+HFD additionally showed significant positive correlations with M4_Synapse_Neuron, M7_MAPK_Metabolism, and M43_Ribonucleoprotein_Binding modules (Figure 4A; Supplementary Table D), whereas other mice (LOAD1+CD, LOAD1+HFD and LOAD2+CD) were positively correlated but below the significance threshold.

**FIGURE 4:**
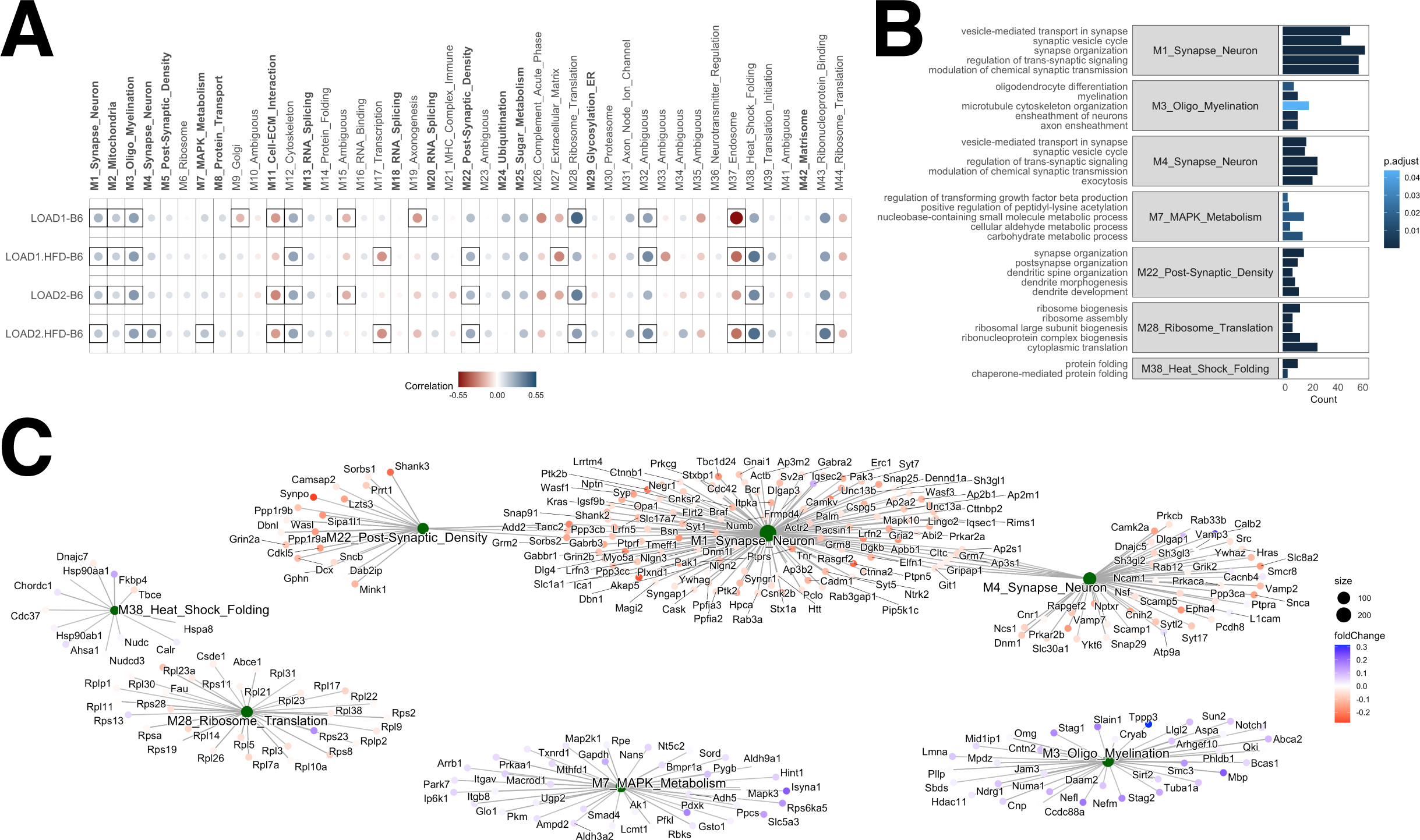
LOAD mice exhibit proteomics changes similar to human LOAD. Correlation coefficients between 18-month-old LOAD mouse models and 44 human proteomics co-expression modules [Johnson et al Ravi’s Ref 9] (A). Modules in bold face were significantly correlated to one or more AD traits. Circles corresponds to positive (blue) and negative (red) Pearson correlation coefficients for protein expression changes in LOAD mice (log fold change of LOAD strains versus B6J) and human disease (log fold change for cases versus controls). Color intensity and size of the circles are proportional to the Pearson correlation coefficient, with significant correlations (p < 0.05) framed. LOAD1 and LOAD2 mice were significantly and positively correlated (p < 0.05) with multiple human AD modules, primarily related to synaptic function. Five top enriched GO terms for proteins with common directional changes for 18-month-old LOAD2 mice on HFD and human AD cases (B). Protein module network with common directional changes for 18-month-old LOAD2 mice on HFD and human proteomics modules (C). Blue (red) nodes correspond to increased (reduced) protein abundance in both 18-month-old LOAD2 HFD mice compared to B6J mice and human AD cases versus controls.

Overall, we detected genetic effects from aged LOAD1 and LOAD2 mice that correlated with multiple human AD proteomics modules that were generally enhanced by exposure to HFD, especially in LOAD2 mice. LOAD2+HFD were positively correlated with ten of the 44 total protein modules, of which five modules (M1_Synapse_Neuron, M3_Oligo_Myelination, M4_Synapse_Neuron, M7_MAPK_Metabolism, and M22_Post-Synaptic_Density) were correlated with AD traits^19^. These modules frequently represented neuronal proteins, distinct from the immune and metabolic signatures observed in transcriptomic analyses.

To better understand the functions of proteins driving the significant positive correlations between LOAD2 mice on HFD and human AD, we isolated the proteins within each module with common directional changes (increased or decreased abundance) for LOAD2+HFD and human AD cases. We performed gene ontology enrichment analysis on these proteins. Proteins that showed mouse-human directional coherence in the M1_Synapse_Neuron module were enriched for biological functions including “synaptic vesicle cycle”, “vesicle-mediated transport in synapse”, and “synapse organization” (Figure 4B; Supplementary Table E). Proteins that showed directional coherence in the M4_Synapse_Neuron module were enriched for “synaptic vesicle cycle” and “exocytosis” biological functions (Figure 4B; Supplementary Table E). In the M22_Post-Synaptic_Density module, coherent mouse-human proteins were enriched for biological functions including “synapse organization”, “postsynapse organization”, and “dendrite development” (Figure 4B; Supplementary Table E). These functions represent the core neuronal processes recapitulated in LOAD2+HFD at the protein level.

Proteins in these synaptic AD protein modules mostly had reduced abundance in LOAD2.HFD mice (logFC < 0) compared to chow-fed B6J controls. This reduced expression of proteins associated with synapse/neuronal functions was similar to human AD cases (Figure 4C), although we note these mice did not exhibit frank neurodegeneration.

For non-synaptic modules, we found coherent proteins in the M3_Oligo_Myelination module were enriched for biological functions including “oligodendrocyte differentiation”, “myelination”, and “microtubule organization” (Figure 4B; Supplementary Table E). Proteins with directional coherence in the M7_MAPK_Metabolism module were enriched for biological functions such as “regulation of transforming growth factor beta production” and “carbohydrate metabolic process” (Figure 4B; Supplementary Table E). In the M28_Ribosome_Translation module, protein abundances mostly increased (logFC > 0) and were enriched for “ribosome biogenesis” and “cytoplasmic translation” (Figure 4B-C; Supplementary Table E). Finally, coherent proteins in the M38_Heat_Shock_Folding module were enriched for “protein folding” and “chaperone-mediated protein folding” biological functions (Figure 4B; Supplementary Table E). Proteins exhibiting directional coherence in the M3_Oligo_Myelination and M7_MAPK_Metabolism modules generally had greater abundances in LOAD2+HFD mice compared to B6J controls and human AD cases, corresponding to increased protein expression associated with ‘MAPK_metabolism’ and ‘Oligo myelination’ (Figure 4C).

### Longitudinal Volumetric Measurements reveals age-dependent changes in brain volume (cohort 2)

Longitudinal volumetric measurements were employed to elucidate age-dependent alterations in brain volume within Cohort 2. Building upon the findings derived from Cohort 1 at JAX, which encompassed indicators such as neuronal cell loss, increased levels of neurofilament light chain (NfL), and molecular manifestations of neuronal cell dysfunction (refer to Figures 1-4), Cohort 2 was established at Indiana University. This cohort consisted of male and female mice designated as LOAD1 and LOAD2, subjected to either a standard control diet (CD) or a high-fat diet (HFD). The primary objective at Indiana University was to assess brain volumes through in vivo magnetic resonance (MR) imaging and to conduct a comprehensive evaluation of LOAD2+HFD mice for plasma and brain biomarkers, with potential implications for preclinical testing.

In vivo MR imaging was conducted on LOAD2 CD and HFD-fed mice at 4, 12, and 18 months of age. The mean whole brain volumes for male LOAD2 CD and HFD mice were 478.56±18.63 and 459.56±6.41 mm^3 at 4 months, 464.53±15.16 and 468.40±9.83 mm^3 at 12 months, and 502.68±13.59 and 470.25±6.37 mm^3 at 18 months, respectively. Correspondingly, the mean whole brain volumes for female LOAD2 CD and HFD mice were 468.19±7.28 and 467.42±6.52 mm^3^ at 4 months, 479.84±12.60 and 483.48±8.20 mm^3^ at 12 months, and 507.29±11.46 mm^3^ and 487.37±11.86 mm^3^ at 18 months.

Statistical analysis revealed a significant reduction in brain volume was observed at 4 and 18 months in LOAD2+HFD male mice (4 months, p=0.0225 and 18 months, p=7.88e-6), while no significant difference was observed at 12 months (p=0.5134). For LOAD2 female mice on a HFD, a significant difference was observed at 18 months (p=0.0186), but no significant disparities at 4 months (p=0.812) or 12 months (p=0.4911).

Among the 165 brain labels analyzed, 45, 57, and 95 brain areas exhibited significant volumetric reductions at 4, 12, and 18 months, respectively, for male mice. Similarly, for female mice, 47, 67, and 51 brain areas displayed significant reductions at 4, 12, and 18 months. Volumetric statistical maps were generated for both male and female LOAD2 CD and HFD mice at the time points, and the significant areas were superimposed onto the T2-weighted template image. Notably, the analyses unveiled a progressive increase in the number of significant areas with advancing age of the mice (Figure 5A).

**FIGURE 5.**
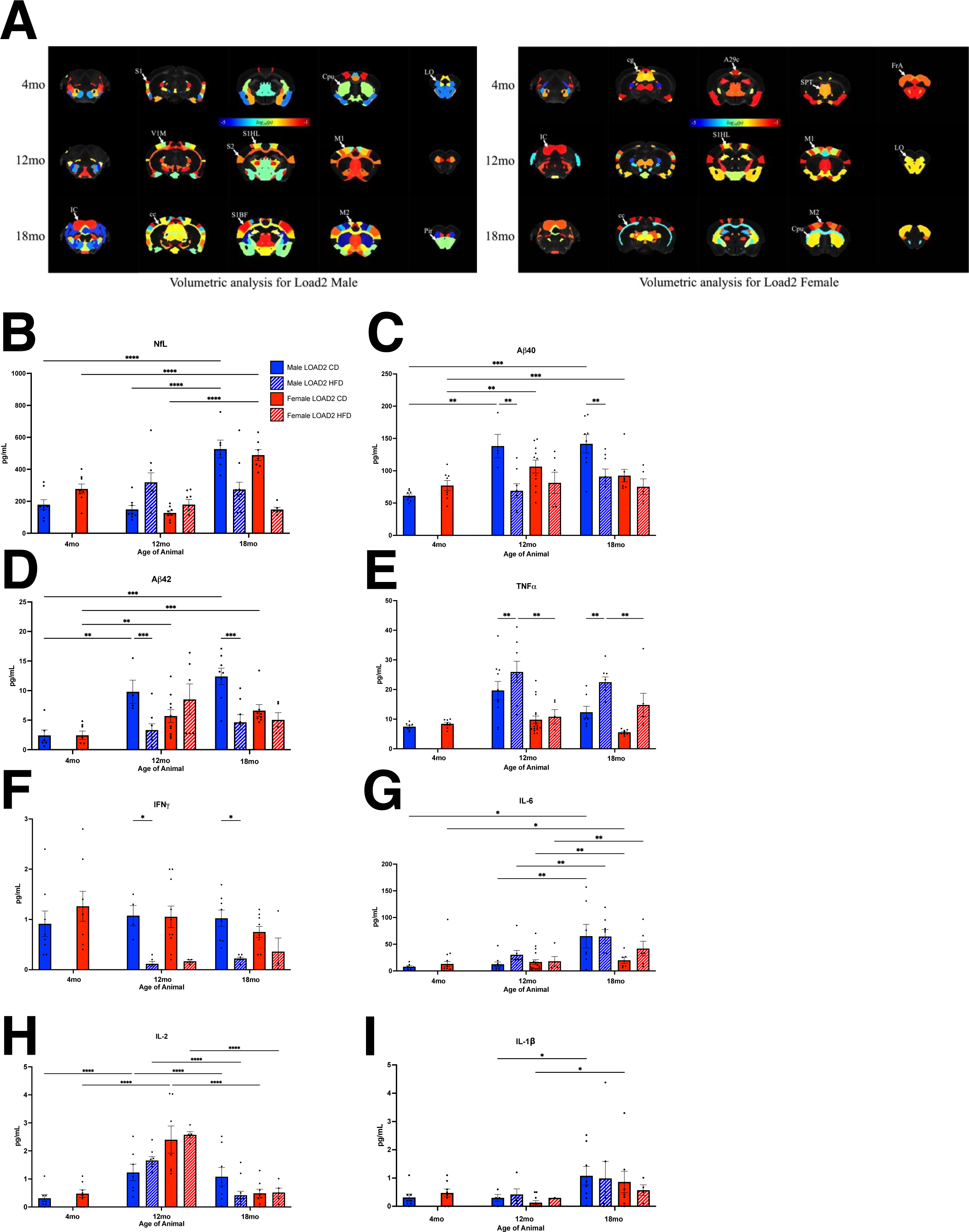
High fat diet reduces brain volume in multiple brain regions and alters plasma biomarkers. Volume statistics maps for LOAD2 male at 4 months, 12 months and 18 months. The significant brain areas were overlaid over gray scale subject template image. (The p-values were converted into logarithmic scale between range ™5 to ™1. S1, Primary Somatosensory Cortex; Cpu, striatum; LO, Lateral Orbital Cortex; V1M, Primary Visual Cortex, Monocular area; S2, Secondary Somatosensory Cortex; S1HL Primary Somatosensory Cortex, Hindlimb; S1BF Primary Somatosensory Cortex, Barrel Field; M1, Primary Motor Cortex; M2 Secondary Motor Cortex; IC Inferior Colliculus; cc Corpus Callosum; Pir, Piriform cortex. Volume statistics maps for Load2 female at 4 months, 12 months and 18 months. The significant brain areas were overlaid over gray scale subject template image. The p-values were converted into logarithmic scale between range ™5 to ™1. A29c, Cingulate Cortex; cg, Cingulum; SPT, Septum; FrA, Frontal Association Area; IC, Inferior Colliculus; S1HL, Primary Somatosensory Cortex, Hindlimb; LO, Lateral Orbital Cortex; M1 Primary Motor Cortex; M2 Secondary Motor Cortex; cc, Corpus Callosum; Cpu, Striatum) (A). At 12 months of age, NfL is increased in high fat diet males, but not in females (B). However, by 18 months of age, aging has a greater effect on NfL, with increases in NfL between 12 and 18 months in both males and female mice. Plasma Aβ40 is reduced in high fat diet males at 12 and 18 months of age but is unchanged in female mice at any timepoint (C). Aβ42 is not altered by a high fat diet (D). TNF-α is increased in male mice on a high fat diet at both 12 and 18 months of age (E). Females have increased TNF-α at 18 months of age, but it does not reach the level of significance. IFNγ is reduced at 12- and 18-month male mice on a high fat diet (F). IL-6 is increased in male mice between 12-18 months of age, but there is no effect of high fat diet (G). In females, IL-6 is not altered. IL-2 increases between 4- 12 months in males and females regardless of diet (H). By 18 months, IL-2 is reduced in both males and females. IL-1β increases with age between 12-18 months regardless of diet (I). *p<0.05, **p<0.01, ***p<0.001, ****P<0.0001.

### Plasma and Brain Cytokines Are Altered by HFD

We next used cohort 2 mice to identify cytokines in the plasma and brain that may be utilized as biomarkers for preclinical testing. Analysis of NfL and cytokines in the plasma revealed an increase in NfL at 12 months in HFD males (Figure 5B). By 18 months, however, the HFD effect was absent, but an increase in NfL was driven by age (Figure 5A). Aβ40 was lower in males fed a HFD at 12 and 18 months, but the same difference was not observed in females; Aβ42 levels were not significant at any time point, likely due to variability among groups, however, there was an overall trend for a reduction in Aβ42 in males fed a HFD at 12 and 18 months (Figure 5 C,D). In HFD animals, we found significant increases in TNFα (Figure 5E) in LOAD2+HFD males at 12 and 18 months, females were approaching a significant increase by 18 months, but did not reach significance (Figure 5F). Additional proinflammatory cytokines were examined (Figure 5 G-I), with reductions in IFNγ observed in HFD males at 12 and 18 months and age-related alteration in IL-6, IL-2 and IL1β. Interestingly, as a confirmation and extension of these data and to demonstrate cross-laboratory replicability, we also observed sustained TNFα levels in HFD mice from Cohorts 4 and 5 (University of Pittsburgh). LOAD2+HFD males from 2 months of age in Cohort 4 maintained significantly higher concentrations of TNFα in their plasma from 7 months of age onward compared to LOAD2 males fed CD, while females on HFD appeared to follow a similar trend as males but only reached significance at 8-8.5 and 15-18 months of age (Figure 8B).

At 18 months of age, the brains from longitudinal cohort 2 were processed for biochemistry. In the brain, female LOAD2+HFD had increased insoluble Aβ42 (Figure 6A), but both males and females on a HFD had reduced insoluble Aβ40 and soluble Aβ40 and 42 (Figure 6 B-D). With regards to proinflammatory cytokines, HFD reduced IL-5 and IL-4 in females (Figure 6 F-G), but increased IL-2, KC-GRO, and IL-12 in both males and females (Figure 6 H-J). Interesting, in the brain (as compared to plasma), TNFα (Figure 6K) remained unchanged. Proinflammatory IL-10 was increased in males and females on a HFD (Figure 6L).

**FIGURE 6:**
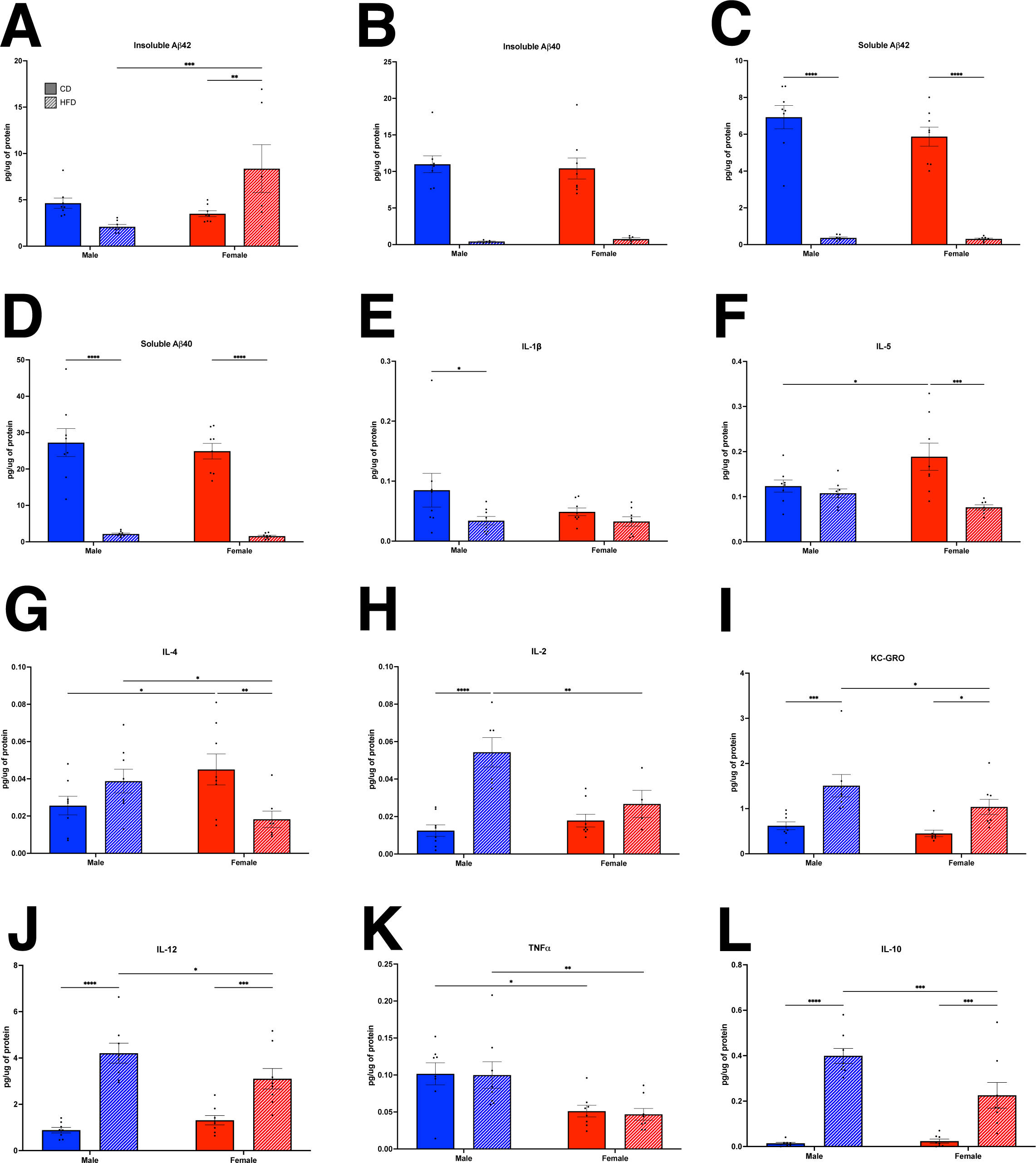
Brain biomarkers in a longitudinal cohort of LOAD2 mice fed a high fat diet. At 18 months, insoluble Aβ42 is increased in female mice on a high-fat diet (A), however, insoluble Aβ40 is decreased in both sexes (B). Soluble Aβ40 (C) and Aβ42 (D) are both reduced at 18 months in high fat diet animals of both sexes. IL-1β is reduced in males on a high fat diet (E) but remains unchanged in females. IL-5 (F) is unchanged in male mice, but is reduced in females on a HFD. IL-4 (G) is also reduced in females fed a HFD. IL-2 (H) is increased in males, but not females. KC-GRO (I) and IL-12 (J) are increased in both males and female mice on a high fat diet. TNF-α (K) remains unchanged on a high fat diet. IL-10 (L) was significantly increased in males and females on HFD. *p<0.05, **p<0.01, ***p<0.001, ****P<0.0001.

### In vivo PET/CT Analysis of Regional Neurovascular Uncoupling (cohort 3)

Cohort 3 mice (female and male LOAD1 and LOAD2 mice on CD and HFD) were established at Indiana University (see **Methods**) to assess neurovascular uncoupling of ^18^F-FDG and ^64^Cu-PTSM, as measurements of cerebral glucose uptake and brain perfusion. Consistent with transcriptomics (Figure 3-4), along with blood and brain cytokines (Figures 5-6), the addition of HFD to 12 mo LOAD1 mice (relative to 12 mo LOAD on CD) resulted in a Type 1 neurovascular uncoupling of perfusion and glycolytic metabolism, which was sexually dimorphic in nature. Across multiple brain regions, denoted by region annotation (Figure 7A), female LOAD1 mice showed a significant reduction in glucose uptake, coupled with a hyperemia of the same brain regions, consistent with a cytokine driven diabetic phenotype. Statistical comparison revealed that dorsal-medial-ventral areas of the Auditory Cortex (AuDMV), Dysgranular Insular Cortex (DI), Lateral Orbital Cortex (LO), Primary Motor (M1) and Secondary Motor (M2) Cortex, Parietal Association Cortex (PtA), Retrosplenial Dysgranular Cortex (RSC), Primary Somatosenory Cortex (S1), and Thalmus (TH) were significantly different (p<0.05, unpaired t-test) relative to the control diet groups in female mice (Figure 7A, top panel). By comparison, regional uncoupling analysis of male LOAD1+HFD only showed significant changes (p<0.05, unpaired t-test) in the Dorsolateral Orbital Cortex (DLO), Primary Motor (M1) and Secondary Motor (M2) Cortices.

**FIGURE 7:**
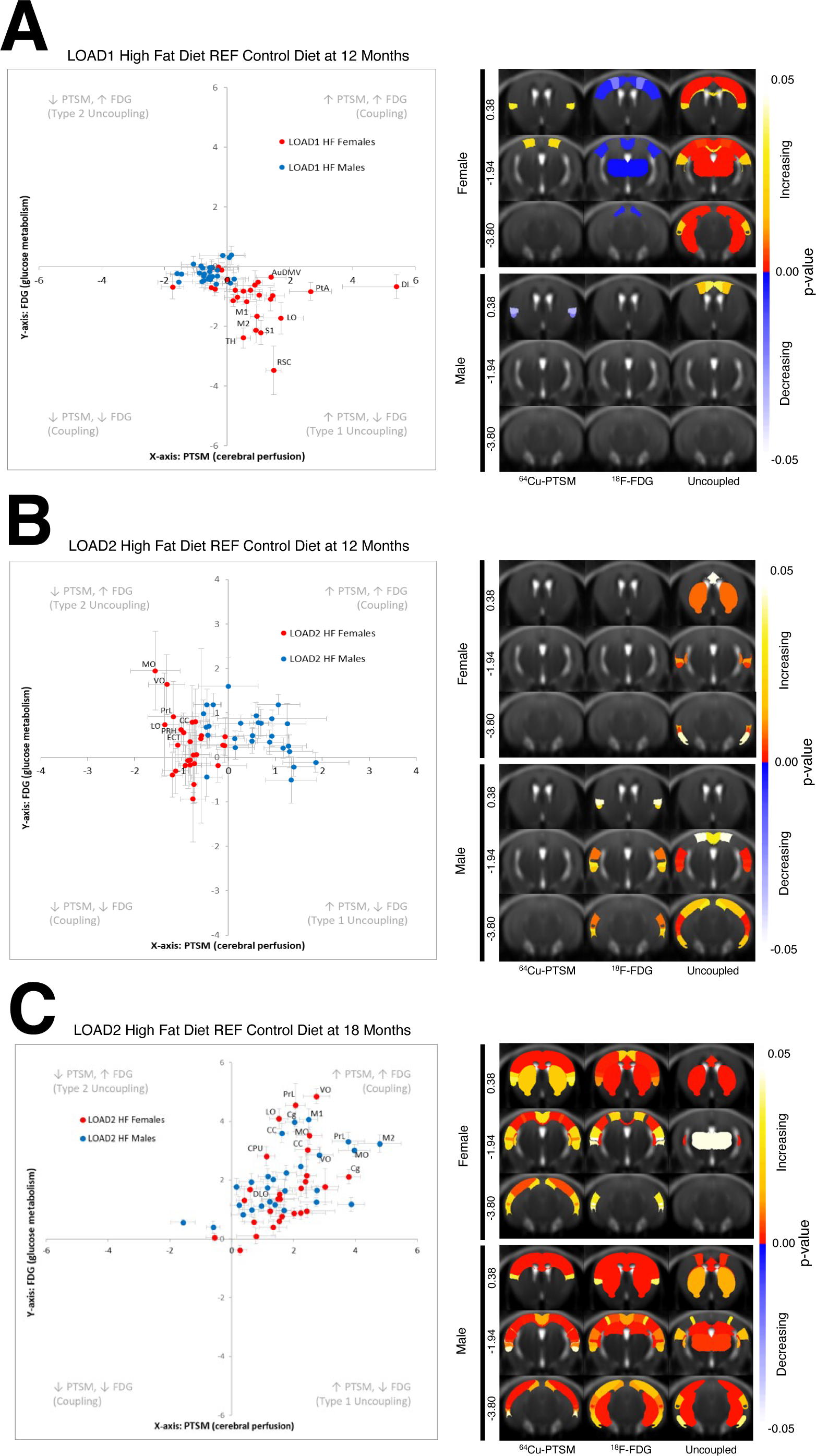
Neurovascular Uncoupling of LOAD1 and LOAD2 mouse models. The degree of neurovascular coordination in LOAD1 (A) and LOAD2 mouse models (B) conditioned on high-fat diet (HFD), we performed uncoupling analysis. (Left) Uncoupling analysis chart in male (blue) and female (red) mice at 12 months, with many brain regions showing significant decreases in metabolism with increases in perfusion. LOAD2 animals aged to 18 months (C) were similarly analyzed. (Upper Right) Female and (Lower Right) Male p- value males showing which regions were significantly different for perfusion, metabolism, and uncoupling.

Importantly, the addition of hAβ onto the LOAD1 background (yielding LOAD2), showed a Type 2 neurovascular uncoupling phenotype when placed on a HFD (relative to LOAD2 on a CD), which was only observed in the female cohort. Unlike LOAD1 mice, female LOAD2 mice on a HFD showed a significant increase in glucose uptake concomitant with regional reductions in tissue perfusion (Figure 7B) and aligned with the transcriptomic network changes (Figure 3B), and blood cytokine levels for TNFα, IL1β and IL6 (Figure 5). Statistical analysis of brain regions revealed that Corpus Callosum (CC), Entorhinal Cortex (ECT), LO, Medial Orbital Cortex (MO), Perirhinal Cortex (PRH), Prelimbic Cortex (PRL), and Ventral Orbital Cortex (VO) all were significantly different (p<0.05, unpaired t-test) from control diet groups (Figure 7A, top panel) in female LOAD2 mice. By contrast, male LOAD2+HFD showed different brain regions, which were uncoupled with treatment, with the AuDMV, Dorsintermed Entorhinal Cortex (DLIVEnt), ECT, PRH, RSC, Temporal Association Cortex (TeA), and Primary and Secondary Visual Cortex (V1V2).

To elucidate the role of aging on gene x environmental effect, we performed neurovascular uncoupling analysis on LOAD2 mice at 18 mo. Unlike LOAD1 and LOAD2 at 12 mos which showed a Type 1 and Type 2 uncoupling phenotype respectively, LOAD2 at 18 mos revealed a hypermetabolic and hyperemic phenotype in both sexes, which resulted in nearly all brain regions increasing in glucose uptake paired with increases in tissue perfusion (Figure 7C). This increase in both perfusion and metabolism at this age, and is consistent with the plasma cytokine data (Figure 8) that show a significant elevation in TNFα, IL-6 and IL-5, which are markers of cell activation and proliferation and are consist with clinical reports of prodromal conversion from healthy controls to MCI.

**FIGURE 8:**
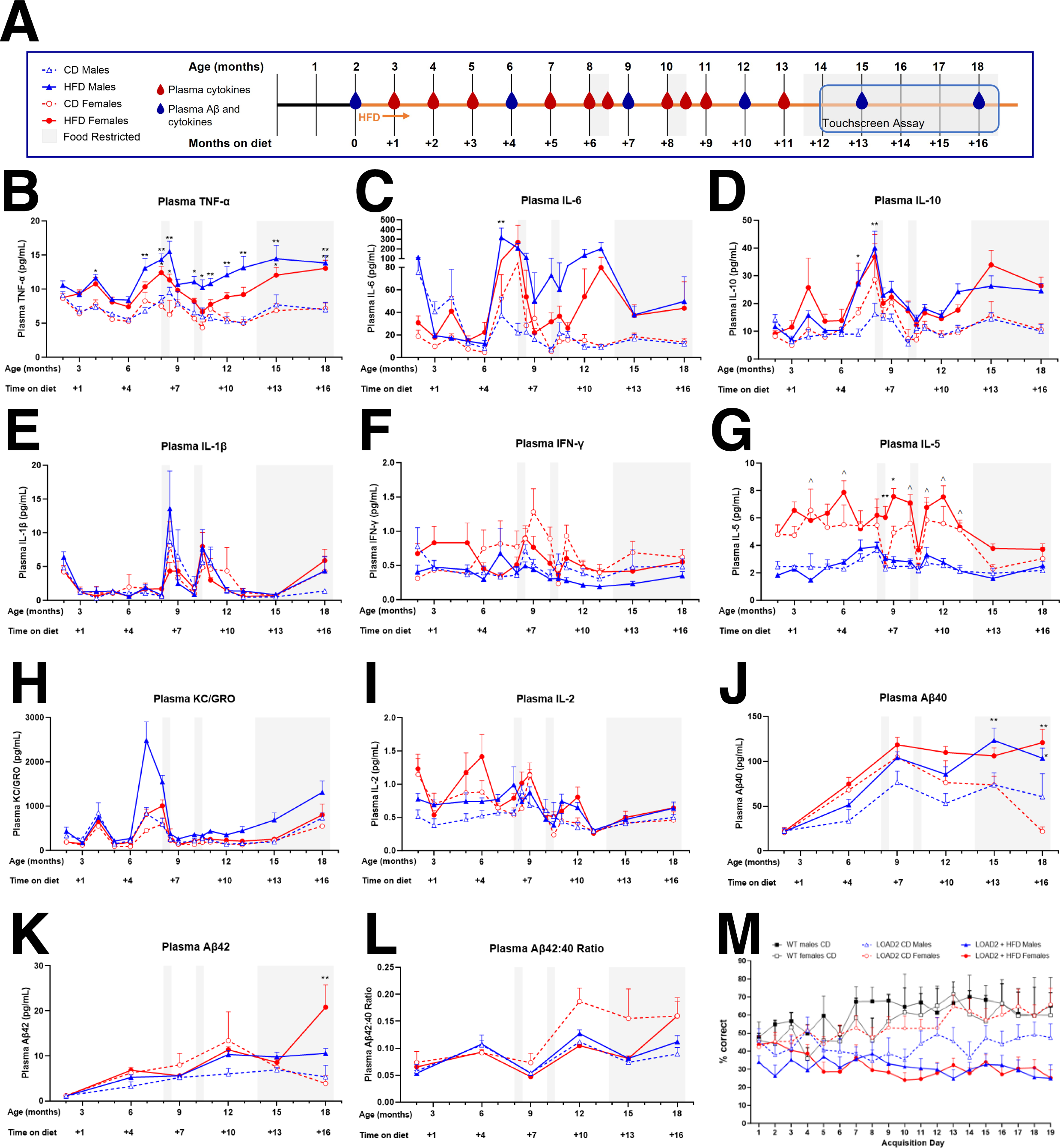
Comprehensive validation of LOAD2 mouse model for preclinical drug testing. As a confirmation and extension of initial characterization data of the LOAD2 mouse model conditioned on high-fat diet (HFD) to serve as a potential model for preclinical testing, independent cohorts were evaluated for disease trajectory of serial plasma biomarkers and cognitive testing. A) Illustration of timeline and procedures; B) plasma TNF-α (pg/mL); C) Plasma IL-6 (pg/mL); D) plasma IL-10 (pg/mL); E. Plasma IL-1β (pg/mL); F) plasma IFNy (pg/mL), G) plasma IL-5 (pg/mL); H) plasma KC-GRO (pg/mL); I) plasma IL-2 (pg/mL); J) plasma Aβ40 (pg/mL); K) plasma Aβ42 (pg/mL); L) calculated Aβ 42:40 ratio in plasma; M) Learning curves of aged (14+ month) LOAD2 mice ± HFD in comparison to age- and sex-matched WT controls during the acquisition phase of the touchscreen cognitive testing battery. Plasma cytokines and plasma Aβ were measured using MesoScale Discovery multiplex ELISA kits (in accordance with the manufacturer’s protocol.

### Evaluation of age-dependent plasma biomarkers and Touchscreen cognitive testing (Cohorts 4 and 5)

To further consider the relevance of the LOAD2 mice as a model for preclinical testing, a group of male and female LOAD2 mice fed a CD or HFD from 2 months of age (cohort 4) and a group of male and female LOAD2 and wildtype littermate controls (WT) fed CD or HFD from 6+ months of age (cohort 5) were established at the University of Pittsburgh. The goal of these cohorts was to track longitudinal biomarker measures for cytokines and Aβ throughout aging, in order to investigate the inflection point at which biomarkers revealed pathological consequences of environment x aging x gene effects, in order to identify a window for therapeutic intervention for future preclinical testing. In addition to biomarker analysis through aging, the cohort were also evaluated for cognitive function assessed by a touchscreen testing battery as well as evaluating the effects of food restriction necessary for touchscreen testing on cytokine levels (Figure 8 and Supplemental Figure 7).

Similar to cohort 2 at IU (Figure 5E), LOAD2 males fed HFD in cohort 4 had increased levels of plasma TNFα at 12 and 18 months of age relative to LOAD2 males fed CD (12 months, p=0.0001 and 18 months, p=0.0002). In fact, analysis of monthly plasma cytokines revealed increased TNFα as early as 4 months of age (p=0.0377) with consistent increases from 7 months of age onward in LOAD2+HFD males compared to LOAD2+CD males (p<0.05, Figure 8B).

The effects of HFD on plasma TNFα was less robust in females, consistent with data from cohort 2 at IU. We observed an overall trend of higher concentrations of plasma TNFα in females fed HFD compared to females fed CD, but those levels were significantly different only at 8-8.5 months (p<0.05) and 15-18 months of age (p=0.0198 at 15 months and p=0.0068 at 18 months; Figure 8A). In addition to TNFα, we also measured 9 other cytokines, 7 of which were above the lower limits of detection (see **Supplemental methods**). Plasma levels of IL-6 (p=0.0022) and IL-10 (p=0.0375) were significantly higher in LOAD2+HFD males compared to LOAD2+CD males at 7 months of age, and levels of IL-10 (p=0.0025) but not IL-6 (p=0.0832) were also higher at 8 months of age in HFD males (Figure 8C, D) prior to the first trial of food restriction that preceded touchscreen testing. Interestingly, plasma levels of IL-5 were significantly higher in LOAD2 females, regardless of diet, compared to males at several time points (p<0.05, Figure 8G) while other plasma cytokines measured – IL-1β, IFNγ, KC-GRO and IL-2 (Figure 8E, F, H, and I) – did not show diet- or sex-related effects in this cohort. In cohort 5, the middle-aged (6- 12 months old) start for HFD (Supplemental Figure 7A) produced a more modest effect on plasma TNFα with LOAD2+HFD males showing higher concentrations of plasma TNFα at 4 months on diet relative to WT males on CD (p=0.0241) and at 5 months on diet relative to LOAD2 and WT males on CD (p=0.0451 and p=0.0034), LOAD2+HFD females showing higher concentrations of TNFα relative to LOAD2+CD females only at 5 months on diet (p=0.0291), and WT males but not females on HFD showing higher concentrations of TNFα relative to WT+CD mice also only at 5 months on diet (p=0.0091 males and p=0.998 females, Supplemental Figure 7B). In all other cytokines measured – IL-6, IL-10, IL-1β, IFNγ, IL-5, KC-GRO and IL-2 (Supplemental Figure 7C-I) – there was no significant effect of diet on LOAD2 or WT males or females, though we did again observe a sex-effect in plasma IL-5 with concentrations being higher in LOAD2 females compared to LOAD2 males regardless of diet (p<0.01 at 2 months on diet and p<0.05 at 3 months on diet, Supplemental Figure 7G).

Concentrations of plasma Aβ40 and 42 were also altered by HFD in cohort 4 (Figure 8J-K) and by genotype but not diet or sex in cohort 5 (Supplemental Figure 7 J,K) consistent with the observation that HFD started earlier in life (cohort 4) produces a more robust phenotype than when initiated mid-life (cohort 5). Male LOAD2 mice +HFD beginning from 2 months of age had increased levels of plasma Aβ40 at 15 (p=0.01) and 18 months of age (p=0.0365) relative to LOAD2 +CD males while female LOAD2+HFD beginning from 2 months of age had increased plasma Aβ40 at 18 months (p<0.0001) relative to LOAD2+CD females (Figure 8J). As demonstrated in Fig S6A LOAD2 mice beginning HFD at 6+ months of age (cohort 5) failed to demonstrate increases in plasma Aβ40 relative to LOAD2+CD (Supplemental Figure 7J). Plasma Aβ42 was significantly increased in LOAD2+HFD females from cohort 4 compared to LOAD2+CD females at 18 months of age (p<0.0001, Figure 8K) and compared to age-matched WT+CD females (p=0.0319, Figure S6K).

While there was robust cross-cohort and cross-laboratory replicability for plasma cytokines irrespective of the different laboratory environments, we observed divergent results for plasma Aβ for cohort 2 (Figure 5C, D) versus cohorts 4 and 5 (Figure 8J, K and Supplemental Figure 7J, K). Two major factors may contribute to these differences beyond different laboratory environments: 1) plasma Aβ was analyzed from anesthetized subjects during terminal procedures for cohort 2 whereas cohorts 4 and 5 were longitudinally sampled in non-anesthetized mice; and 2) Cohorts 4 and 5 were subjected to periods of food restriction which was not a factor for cohort 2. It is well documented that variations in plasma Aβ are influenced by environmental factors including stress^20^. Despite the cross-lab variability for Aβ, irrespective of transient stressors, TNFα levels were sustained consistently across cohorts and align with transcriptomic and proteomic data which demonstrates the robustness of the LOAD2 x age x HFD model for preclinical studies of therapeutic interventions independent of amyloid.

We assessed the cognitive function of both cohort 4 and 5 using a touchscreen assay battery. All subjects demonstrated the ability to associate the response (nosespoke touch) with the presentation of a reward as measured by the ability to meet a priori criterion of 2 consecutive sessions of 30 rewards earned within 45 minutes. Interestingly, despite similar food restriction across cohorts, there was a significant effect of diet with either WT or LOAD2 mice regardless of HFD initiation (at 2 months or at 6+ months) failing to acquire the task as measured by % accuracy not exceeding chance levels (50%). LOAD2+HFD mice required a greater number of days to meet criteria relative to LOAD2 mice on control diet [One way ANOVA [F (3, 24) = 5.290; P=0.0061; Figure 8M]. Within sex analysis revealed a statistically significant increase in females reared on HFD (T-test; p<0.05) and a modest non-significant increase in males reared on HFD (T-test; p=0.38). Within control diet groups, there was no effect of genotype or sex (p>0.05). During the punish incorrect phase, 100% of CD mice, irrespective of genotype met criteria however only 27% of LOAD2+HFD male mice met a priori completion criteria for this task while 0% of female LOAD2+HFD mice met criteria. Analysis of accuracy levels during initial acquisition of the task prior to subjects advancing to the location discrimination phase (Training Days 1-18) revealed a statistically significant impairment in HFD treated subjects relative to genotype and age- and sex-matched non-HFD controls as measured by two-way repeated measures ANOVA: [F (5, 33) = 9.047; p<0.001; Figure 8M]. Analysis of WT and LOAD2 mice exposed to HFD at middle-age (6+ months age) revealed a less aggressive phenotype as indicated by lower plasma cytokine levels and lower plasma Aβ40 and Aβ42 when compared with plasma levels from mice exposed to HFD at 2 months of age (Figure 8). **Supplemental Figure 7M** illustrates performance of aged (18+ month) LOAD2 mice on CD (purple diamonds) and age- and sex-matched C57BL/6J (WT) mice on control diet (n=3-5 per genotype) on the LD task for both large (easy) and small (hard) separation conditions (mean ± s.e.m.). As expected, the increase in task difficulty was significant across genotypes as measured by increased number of trials required to reach criterion for easy relative to hard [F (1, 12) = 5.156; P=0.04]. While there was a modest increase in the number of trials to reach criterion in LOAD2 mice relative to WT the effect of genotype was not significant [F (1, 12) = 0.4134; P=0.53].

## Discussion

Despite the recent approval of anti-amyloid therapies, there remains a need for improved therapies for Alzheimer’s disease. A key component of therapeutic testing is the development of mouse models that recapitulate the complexity of LOAD. Here we introduce LOAD2 mice – triple homozygous for *APOE4*, *Trem2*R47H* and hAβ. Although they lack amyloid and TAU pathologies, in combination with a HFD, LOAD2 mice show age-dependent development of key aspects of LOAD. Specifically, at 18 months of age, LOAD2+HFD, show loss of neurons in the cortex, increased insoluble Aβ42, brain region-specific volumetric changes, neurovascular uncoupling, and cognitive deficits, consistent with the prodromal stages of AD. Interestingly, these changes correlated with elevated plasma NfL and cytokine levels – clinically relevant biomarkers associated with LOAD and emphasize the robustness of modeling genetics x age x environment.

Neuronal cell number was evaluated by counting NeuN+DAPI+ cells in cortex and hippocampus and revealed a modest reduction in female LOAD2+HFD mice (Figure 2). Coupled with transcriptomic and proteomic signatures (Figures 3 and 4), neurovascular uncoupling (Figure 7), and cognitive deficits (Figure 8), these data suggest circuit dysfunction in LOAD2+HFD mice. Specific circuits are differentially susceptible to aging and/or AD and so neuronal cell loss may be a result of sporadic loss of neurons or loss of neurons within a specific circuity^21-24^. The mechanism by which neurons die in LOAD2+HFD mice is still to be determined. One recent study used a chimeric model system to show that human neurons in the mouse brains exposed to amyloid died by necroptosis^25^, although mouse neurons did not. Evaluating positive markers of programmed cell death (TUNEL, CASPASE3, MEG3) would provide insight into mechanisms driving reduced cortical neurons in female LOAD2+HFD mice. It is also expected that synaptic changes/loss would precede neuronal cell loss and further work is needed to determine this. Also, myelin integrity was not evaluated and may be disrupted in LOAD2 mice.

As observed in human AD^19^, we detected distinct proteomic and transcriptomic signatures in our LOAD mouse models. Transcriptomic signals tended to represent immune, vascular, and lipid metabolism (Figure 3), whereas proteomic signatures were focused in synaptic and myelination modules (Figure 4). The advent of proteomic technologies that commonly quantify around 10,000 proteins, such as the TMT approach used here, now enable deep characterization to reveal such differences. We also note that AD-relevant transcriptomic changes tended to be driven by age and diet (Figure 3A) while proteomic changes were primarily due to APOE4 and *Trem2*R47H* genetics and somewhat exacerbated by diet. These results demonstrate the importance of multi-omic analyses in fully characterizing causal factors (*e.g.,* genetics, diet, age) and affected processes (*e.g.,* synaptic, immunological) in translational research.

Data-driven analysis of transcriptomes through gene co-expression networks revealed two modules, denoted turquoise and lightyellow, that highlighted a molecular separation between normal age-related changes and changes related to AD-relevant biomarkers (Figure 3). While both modules were enriched for immune response, this enrichment was more significant in the turquoise (p = 1.8 x 10^−49^) than the lightyellow (p = 3.7 x 10^−3^) module (Supplementary Table B3). This suggests a more focused immune component in the lightyellow module, led by cytokine signaling (Figure 3C) and often co-annotated to lipid metabolism (Figure 3D). Furthermore, the lightyellow module was much more correlated with circulating cytokines and NfL, while the turquoise module was linked to IBA1 and GFAP markers (Figure 3B). These results suggest a signature for AD-related transcriptomic changes (the lightyellow module) that is distinct from usual brain aging (the turquoise module) and driven by diet in LOAD mice.

Though no deficits in hippocampal spatial working memory as measured by the spontaneous alternation task were observed in LOAD2+HFD mice (Figure 1), across two separate cohorts, aged mice reared on HFD either from 2 months of age or from 6+ months of age failed to meet acquisition criteria as measured by % correct responses less than 50% (chance levels) indicative of a learning impairment (Figure 8 and Supplemental Figure 7). The lack of ability to learn the task is unlikely to be explained by food motivation, as both HFD and CD groups were sufficiently restricted, and all groups independent of diet or genotype acquired initial touch-reward association. This indicates that failure of HFD mice to accurately perform the task is more likely a feature of impaired learning due to the combination of advanced age x environmental risk (e.g. HFD), than reward salience. Further studies may be required to investigate methods for enhancing motivation for rewards in mice reared on HFD which may include more aggressive water restriction protocols as HFD treated mice even in the presence of strawberry-milk shake reinforcer also have issues with touchscreen tasks (personal communication with Dr. Lisa Saksida, also see^26^). Regardless, for aged WT and LOAD2 mice on control diet, results from daily assessments on task acquisition reveal modest cognitive impairments in LOAD2 relative to WT. While modest, these cognitive deficits are further strengthened by proteomic analysis of LOAD2 mice revealing alterations in synaptic signaling. While we were not able to conduct additional cognitive tests in this advanced aged cohort due to attrition and eventual mortality, as subjects aged to 24 months by the conclusion of testing, these data indicate that translational touchscreen tests may be more sensitive for detecting more specific cognitive domains than traditional single day behavioral tests in mice that have been historically used for assessing cognition.

Commensurate with the cytokines and multi-omic (Figure 3B) associations, cerebral perfusion and metabolism via uncoupling analysis revealed a sexually dimorphic dysregulation with age and genotype (Figure 7,B). Importantly, the addition of a HFD in LOAD1 mice, resulted in Type 1 uncoupling (i.e. reductions in glycolysis and compensatory hyperemia), consistent with a cytokine^27-30^ driven down regulation of insulin receptors^28^, which have been shown to result in a reduction in neuronal glucose uptake via GLUT transporters^29-31^. Importantly, these data closely parallel the Type 2 diabetic phenotype^32, 33^ with reactive hyperemia via activation of eNOS^34^.

To further explore the impact of environment on gene and sex, analysis of neurovascular uncoupling was performed to assess the degree of regional metabolic dysregulation in LOAD2 mice. Unlike the base model, LOAD2+HFD at 12 mos resulted in a Type 2 neurovascular uncoupled phenotype (i.e. increased glycolysis and reduced perfusion) which was only observed in female mice. These data align with cytokines (i.e. TNFα, IL-2) and immunopathology changes (Figure 2, 5) at this age, and are consistent with previous reports of cytokine driven astrocytic proliferation and GLUT1 expression^35^. By contrast, LOAD2+HFD mice at 18 mos, resulted in whole brain increases in perfusion and metabolism. Importantly, these changes were observed in both sexes, and was consistent with reports of prodromal hyper-metabolism^36-39^and hyperemia^40^observed in clinical patients, suggesting that this model recapitulates the earliest manifestations of LOAD.

Collectively, data presented here suggest that LOAD2+HFD mice, particularly are presenting with early stages of LOAD by 18 months of age. Further work is needed to develop models that present a wider range of LOAD pathologies. We were unable to determine whether aging LOAD2+HFD longer would have enhanced their LOAD-relevant phenotypes. This is because LOAD2+HFD mice showed an increased incidence of tumors when aged beyond 18 months of age. Control mice, including WT mice, showed similar phenotypes, suggesting this resulted from chronic consumption of HFD (from 2 months). Interestingly, LOAD2+CD mice aged to 24 months or LOAD2+HFD after 12 months of age did not show as severe phenotypes as 18 months old LOAD2H+HFD from 2 months of age, highlighting the significance of environmental influence of chronic exposure to HFD. Several additional strategies are being tested to develop improved LOAD models. Alternative diets, e.g., a milder western diet, are being tested that may recapitulate LOAD2-relevant phenotypes of a HFD without the presence of age-related tumors and attrition. We are also evaluating additional genetic risk factors on the LOAD2 background that impact lipid metabolism (*Abca7*A1527G*), neuroinflammation (*Plcg2*M28L*) and vascular health/metabolism (*Mthfr*677C>T*). Perturbing these specific pathways may exacerbate the effects of *APOE4*, Trem2*R47H and hAβ in LOAD2. Tau pathology was not evaluated in LOAD2 mice and was not expected due to the lack of a humanized *MAPT* gene. Strains where the mouse *Mapt* gene has been replaced by human *MAPT* are now available and therefore, mouse models carrying combinations of *hAβ*, *hMAPT*, genetic risk factors (e.g., *APOE4*) are now being created and will be exposed to environmental risk factors such as HFD and toxic metals to evaluate their potential as preclinical models of AD.

In conclusion, the interaction of genetic risk and aging leads to a phenotype worsened by environmental factors, mirroring the risk for LOAD. This includes a pronounced neuroinflammation phenotype with cognitive impairment, unrelated to amyloid accumulation. Multi-omic analysis identified molecular signatures, such as synaptic signaling deficits, aligning with the observed cognitive impairment. The LOAD2 model, characterized by translational blood and imaging biomarkers, emerges as a crucial tool for preclinical drug testing in ADRD patients without prominent amyloid, potentially offering a more representative model of sporadic LOAD.

## Acknowledgments

The results published here are in whole or in part based on data obtained from the AD Knowledge Portal (https://adknowledgeportal.org). Study data were provided by the Rush Alzheimer’s Disease Center, Rush University Medical Center, Chicago. Data collection was supported through funding by NIA grants P30AG10161 (ROS), R01AG15819 (ROSMAP; genomics and RNAseq), R01AG17917 (MAP), R01AG36836 (RNAseq), the Illinois Department of Public Health (ROSMAP), and the Translational Genomics Research Institute (genomic). Additional phenotypic data can be requested at www.radc.rush.edu. Mount Sinai Brain Bank data were generated from postmortem brain tissue collected through the Mount Sinai VA Medical Center Brain Bank and were provided by Dr. Eric Schadt from Mount Sinai School of Medicine. The Mayo RNAseq study data was led by Nilüfer Ertekin-Taner, Mayo Clinic, Jacksonville, FL as part of the multi-PI U01 AG046139 (MPIs Golde, Ertekin-Taner, Younkin, Price). Samples were provided from the following sources: The Mayo Clinic Brain Bank. Data collection was supported through funding by NIA grants P50 AG016574, R01 AG032990, U01 AG046139, R01 AG018023, U01 AG006576, U01 AG006786, R01 AG025711, R01 AG017216, R01 AG003949, NINDS grant R01 NS080820, CurePSP Foundation, and support from Mayo Foundation. Study data includes samples collected through the Sun Health Research Institute Brain and Body Donation Program of Sun City, Arizona. The Brain and Body Donation Program was supported by the National Institute of Neurological Disorders and Stroke (U24 NS072026 National Brain and Tissue Resource for Parkinson’s Disease and Related Disorders), the National Institute on Aging (P30 AG19610 Arizona Alzheimer’s Disease Core Center), the Arizona Department of Health Services (contract 211002, Arizona Alzheimer’s Research Center), the Arizona Biomedical Research Commission (contracts 4001, 0011, 05-901 and 1001 to the Arizona Parkinson’s Disease Consortium), and the Michael J. Fox Foundation for Parkinson’s Research.

The authors are grateful for the technical support of the IUSM Biomarker Core.

The authors would like the thank the Neurobehavioral Phenotyping core at Jackson Laboratory for their assistance and support during the *in vivo* testing of these animals.

The authors are grateful for the technical support of the University of Pittsburgh Preclinical Phenotyping Core facility as well as the following research staff: Gabi Little, Jason Hart, Aman Reddy, Umesh Nepali, Jenny Willis, and Amber Sanders

## Data Availability Statement

The MODEL-AD data sets are available via the AD Knowledge Portal (https://adknowledgeportal.org). The AD Knowledge Portal is a platform for accessing data, analyses, and tools generated by the Accelerating Medicines Partnership (AMP-AD) Target Discovery Program and other National Institute on Aging (NIA)-supported programs to enable open-science practices and accelerate translational learning. The data, analyses and tools are shared early in the research cycle without a publication embargo on secondary use. Data is available for general research use according to the following requirements for data access and data attribution (https://adknowledgeportal.org/DataAccess/Instructions). For access to content described in this manuscript see: https://doi.org/10.7303/syn53128146

## Conflicts of Interest

The authors declare that this research was conducted in the absence of any commercial or financial relationships that could be construed as a potential conflict of interest.

## Funding Sources

The MODEL-AD Center was supported through funding by NIA grant U54AG054345. GH was supported by the Diana Davis Spencer Endowed Chair Research and GC was supported by the Bernard and Lusia Milch Endowed Chair. NW was supported by NINDS grant R01NS125020.

## Consent Statement

Due to the nature of this study, consent for human subjects was not needed.

## Figure Legends

**SUPPLEMENTAL FIGURE 1:**
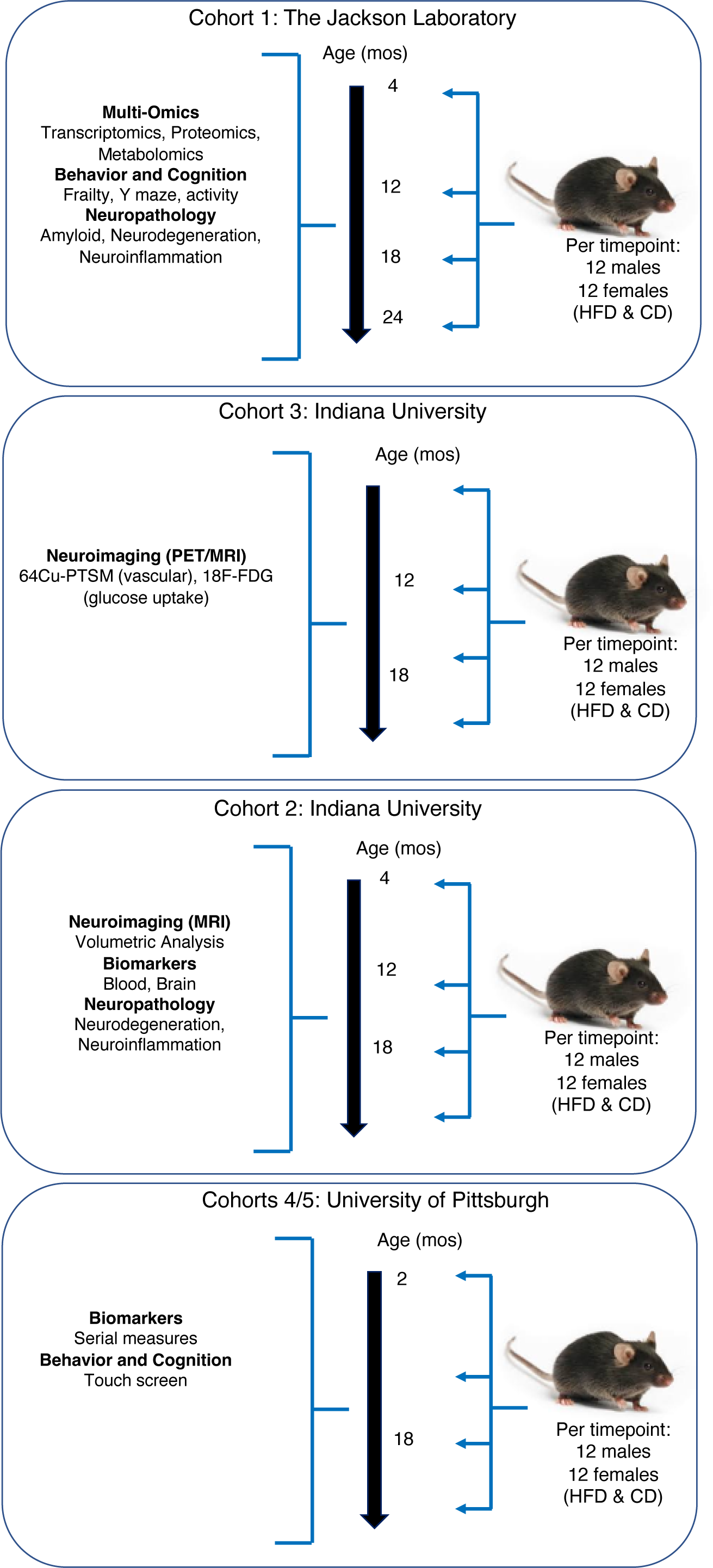
Study design. This was a multi-site, multi-cohort study. At each site, mice were provided a high fat diet or control diet starting at 2 months of age. At the Jackson Laboratory, the 18-month cohort only received a control diet. In cohort 1, four cohorts of mice were aged to 4, 12, 18, 24 months of age. Multiple ‘omics studies, behavior and cognition and neuropathology studies were completed. In cohort 2, at Indiana University, one cohort of mice were age to 18 months and plasma was collected at 4, 12 and 18 months for biomarker analysis. At each timepoint, mice had MRIs completed for volumetric analysis. Brains were collected at the termination of the study for neurodegeneration and neuroinflammation studies. In cohort 3 at Indiana University, two cohorts of mice were aged to 12 and 18 months and underwent PET/CT studies for two tracers, FDG and PTSM. In cohort 4 at University of Pittsburgh, mice were aged to 18 months, undergoing serial blood draws monthly and completing touch screen testing.

**Supplemental Figure 2:**
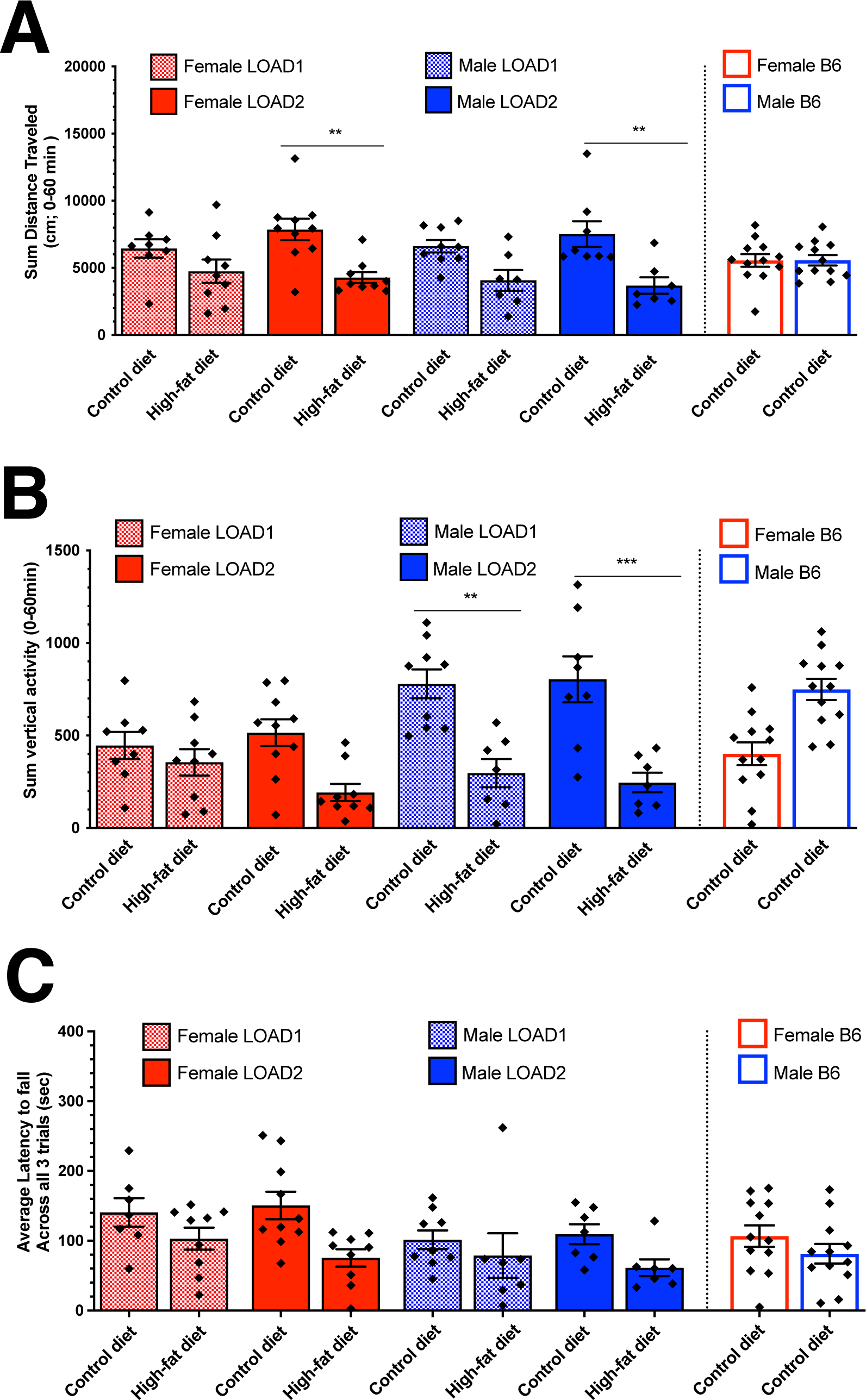
Longitudinal behavioral phenotyping of mice on high-fat diet. Males and females, of LOAD1 and LOAD2 genotypes, fed either control diet (CD) or high-fat diet (HFD) beginning at 2-months of age until 18-months of age were subjected to open field assay, measuring animal movements by way of total distance traveled (A) and total vertical activity (B) during 60 minute observation testing period. Rotarod assay measured the latency to fall times, over three consecutive trials, as a measure of motor coordination (C). (Three-way ANOVA [sex, genotype, diet effects]; *=p<0.05)

**Supplemental Figure 3:**
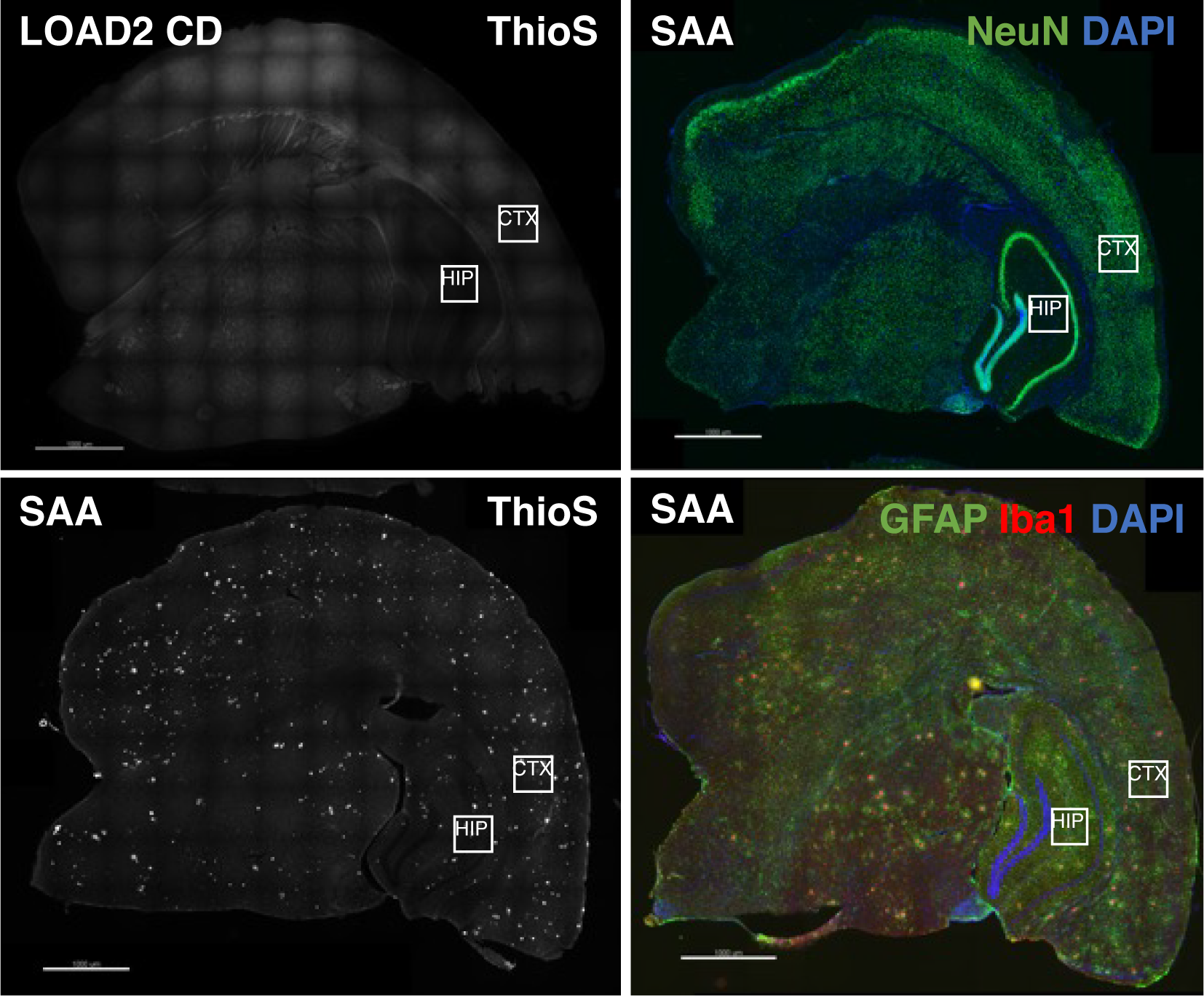
Immunohistochemistry analysis. Representative, collated images of whole hemisphere, coronal sections displaying both cortical and hippocampal regions of interest, used for glia cell density measurements and neuropathological assessment of brain tissue from cohort 1 (JAX). Brains are from 12mo females from either LOAD2 (control diet) and B6J.*APP^SAA^* strains. NeuN=neuronal marker; ThioS=amyloid plaques; GFAP=astrocyte marker; IBA1=microglial marker. Scale bar equals 1,000μm (1mm).

**Supplemental Figure 4:**
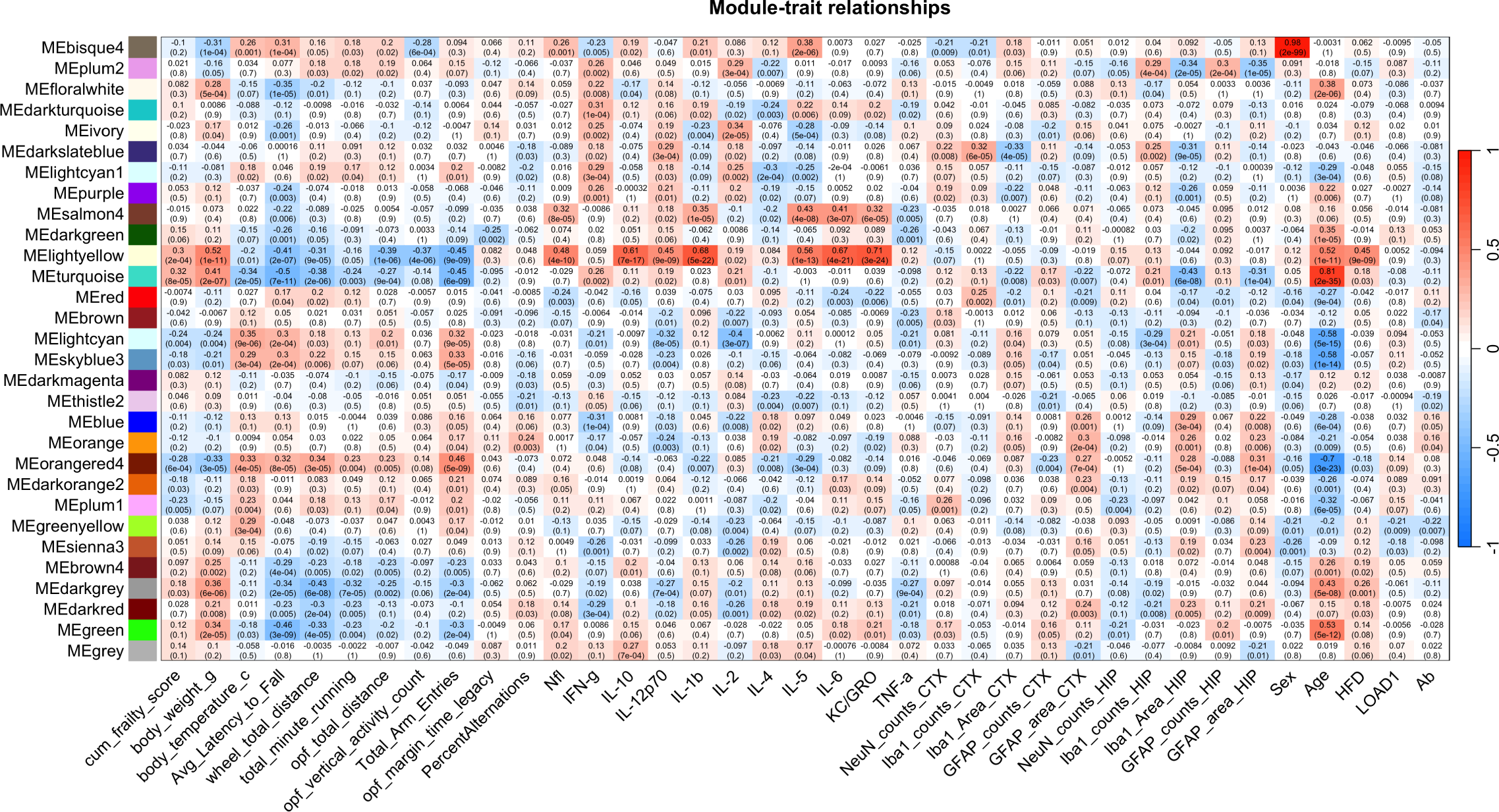
Mouse modules and trait relationships. Transcriptome module correlations and FDR values for module eigengenes and age, sex, diet, genotype, and measured behavioral assays, body weights, NfL, cytokine levels, and GFAP, IBA1, and NeuN cell counts.

**Supplemental Figure 5:**
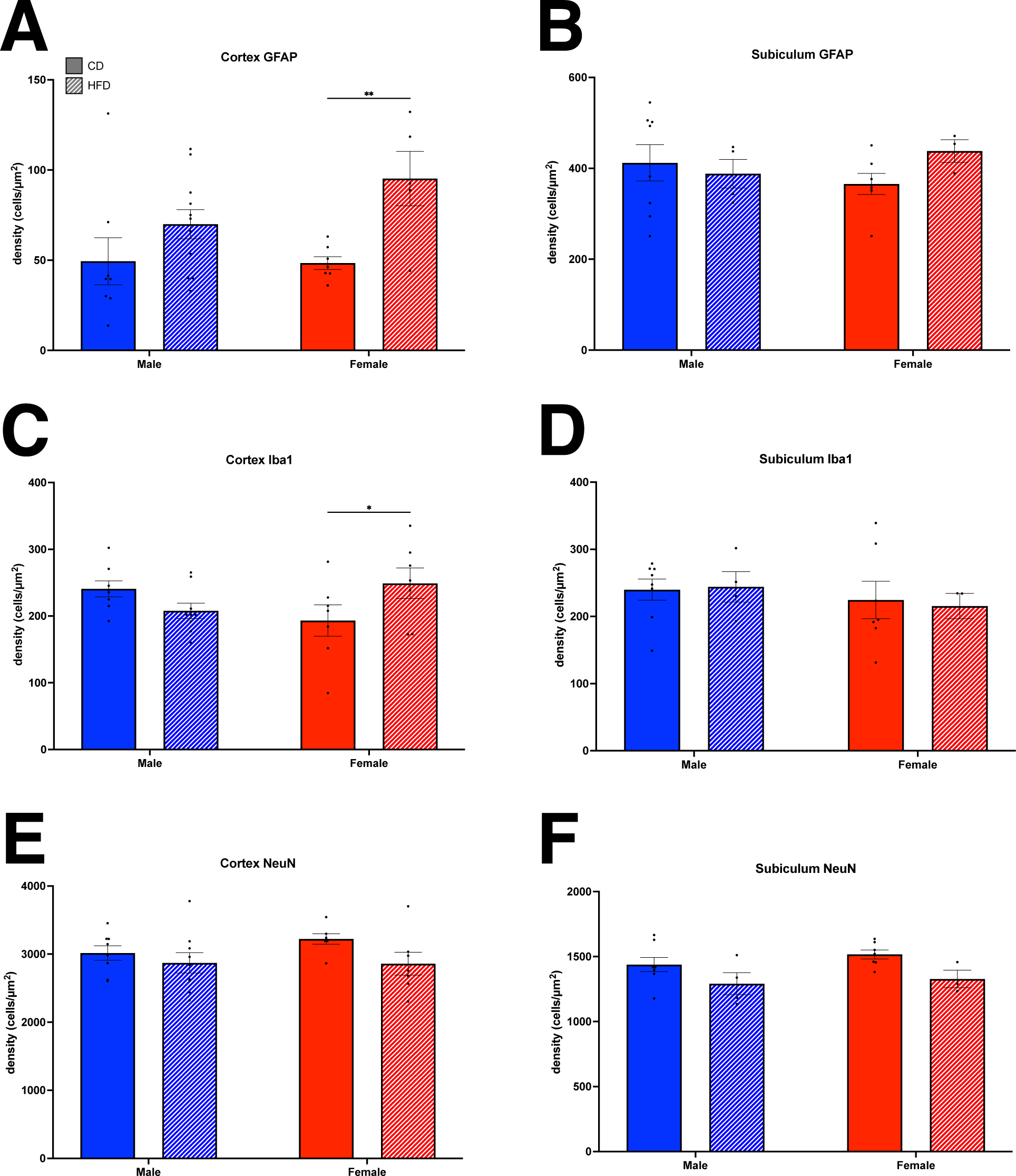
Neurodegeneration and gliosis in the cortex and subiculum. At 18 months old (Indiana University cohort 2), GFAP was used to label reactive astrocytes, which were increased in female mice on a high fat diet in the cortex. IBA1 was utilized to study microglia, which were increased in females on a high fat diet. Neurons (labeled with NeuN) were not changed in the cortex or subiculum.

**Supplemental Figure 6:**
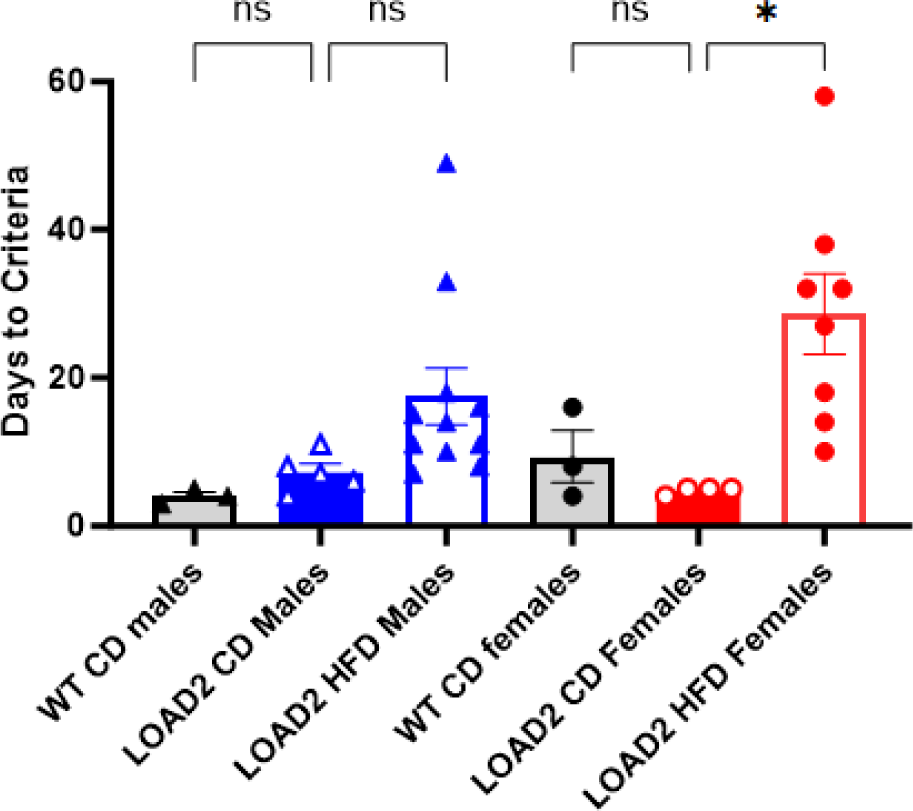
Touchscreen task acquisition is impaired in mice with genetic + environmental risk for AD. Initial touch-reward associations in LOAD2 mice + high-fat diet (HFD) in comparison to LOAD2 and WT mice maintained on a control diet (CD). Data are analyzed as number of days required for individuals to meet a priori task completion criteria.

**Supplemental Figure 7:**
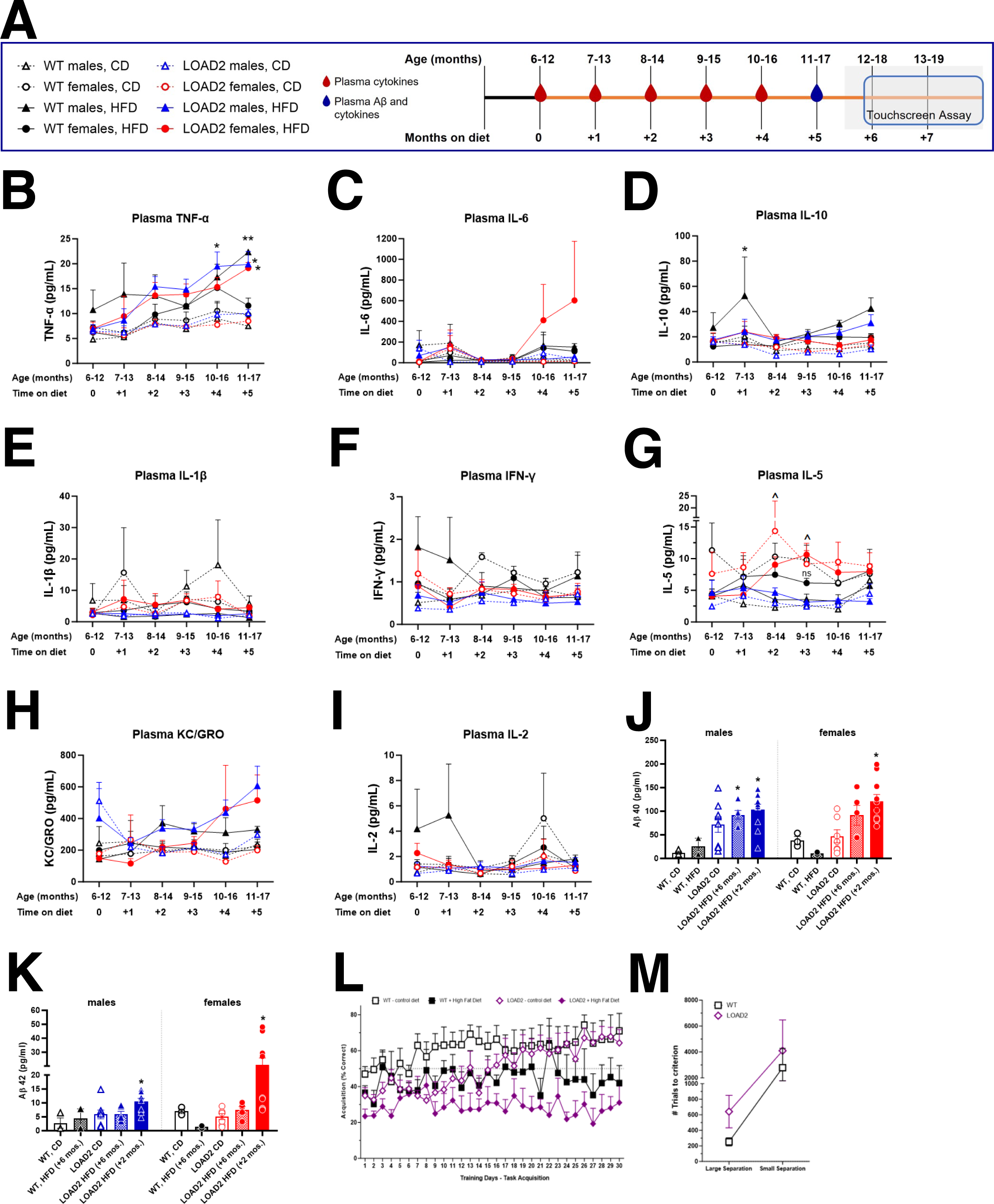
Evaluation of disease trajectory of LOAD2 mice conditioned on a High Fat Diet beginning mid-life at 6 months of age. In comparison to LOAD2 mice conditioned on HFD from 2 months of age, starting HFD after 6 months of age results in a milder phenotype (see Figure 8 for comparison). Chronic HFD exposure produces sustained TNF-α levels in male and female LOAD2 mice relative to WT or LOAD2 mice maintained on control diet (CD); consistent with HFD exposure from adolescence (see Figure 8). Other plasma biomarkers were not significantly elevated and sustained relative to mice on CD. A) Illustration of timeline and procedures; B) plasma TNF-α (pg/mL); C) Plasma IL-6 (pg/mL); D) plasma IL-10 (pg/mL); E. Plasma IL-1β (pg/mL); F) plasma IFNy (pg/mL), G) plasma IL-5 (pg/mL); H) plasma KC-GRO (pg/mL); I) plasma IL-2 (pg/mL); J) plasma Aβ40 (pg/mL) at 12-18 months of age; K) plasma Aβ42 (pg/mL) at 12-18 months of age. Plasma cytokines and plasma Aβ were measured using MesoScale Discovery multiplex ELISA kits in accordance with the manufacturer’s protocol. (\**P*<0.05 and \*\**P*<0.01 between diet groups; ^*P*<0.05 between sexes). One way ANOVA within sex was used to analyze Aβ40 and Aβ42 in 12–18-month aged mice across treatment groups. L) Touchscreen acquisition task data demonstrating impairment of HFD treatment independent of genotype in aged (18+month LOAD2 and WT mice). Both WT+HFD and LOAD2+HFD to fail to learn the task as measured by % accuracy less than chance levels (<50%) (n=3-5 per genotype/diet; combined sexes). Only CD treated WT and LOAD2 mice met acquisition criteria and advanced to assessments of pattern separation as measured by the spatial-location discrimination task. M.) Modest impairments in pattern separation of aged LOAD2 mice (18+ month) relative to age-matched WT (n=3-5 per genotype; sexes combined). Data for touchscreen are presented as mean ± sem.

## SUPPLEMENTARY TABLE A: Differentially expressed genes in mouse models

Differentially expressed genes (DEGs) in mouse strains. Sheet1: DEGs observed in LOAD2 compared to age and sex-matched LOAD1 mice at all ages for both sexes. Sheet 2: DEGs observed in 18-month-old LOAD2 and LOAD1 mice on chow diet and HFD compared to age and sex-matched C57BL/6J mice on chow diet. Sheet 3: DEGs observed in HFD mice compared to sex and strain-matched mice on chow diet at 18 months.

## SUPPLEMENTARY TABLE B: Enriched KEGG pathways in mouse models

Sheet 1: enriched KEGG pathways for the significantly up and down-regulated genes in LOAD2 mouse strains compared to age and sex-matched LOAD1 mice at all ages for both sexes. Sheet 2: enriched KEGG pathways in the significantly up and down-regulated genes in 18-month-old LOAD2 and LOAD1 mice on chow diet and HFD compared to age and sex-matched C57BL/6J mice on chow diet. Sheet 3: enriched KEGG pathways in the significantly up and down-regulated genes in HFD mice compared to sex and strain-matched mice on chow diet at 18 months.

## SUPPLEMENTARY TABLE B2: Mouse modules of co-expressed genes

Genes in each of 30 mouse modules of co-expressed genes identified through WGCNA of mouse transcriptomics data.

## SUPPLEMENTARY TABLE B3: Enrichment of AD biological domains in mouse modules

Enrichment of AD biological domains and member Gene Ontology terms in each mouse module of co-expressed genes.

## SUPPLEMENTARY TABLE C: Differentially expressed proteins in mouse models

Differentially expressed proteins in 18-month-old LOAD1 and LOAD2 mice on chow and HFD compared to age-matched C57BL/6J mice on chow diet.

## SUPPLEMENTARY TABLE D: Correlations between changes in mouse models and case-control changes in 44 human protein co-expression modules

Pearson correlation coefficients and corresponding p-values for the correlation between protein expression changes in LOAD1 and LOAD2 mice relative to B6J mice at 18 months with changes observed in human AD subjects versus controls for each human protein module, as shown in Figure 4A.

## SUPPLEMENTARY TABLE E: Enriched Gene Ontology terms in proteins driving the significant positive correlations between LOAD2 mice on HFD and human AD proteomics modules

Enriched GO terms in proteins with common directional changes for 18-month-old LOAD2 mice on HFD and human AD cases, as shown in Figure 4.

## Supplemental Methods

### Cohort 1 – The Jackson Laboratory

#### Breeding and Husbandry

All mice were housed 5 per cage with SaniChip bedding and initially provided LabDiet® 5K52/5K67 (6% fat; control diet, CD). The mice were kept on a 12-hour light/dark schedule with the lights on from 7:00 a.m. to 7:00 p.m. daily. The animals were ear-punched for identification purposes and subsequently microchipped at the tail base using a p-chip system (PharmaSeq).

#### Behavioral testing

Behavioral tests were conducted as previously reported in the following order with at minimum a 1-2-day rest period between tests: Frailty assessment with core body temperature recording, open field test, spontaneous alternation, rotarod, and wheel running activity. On each test day, subjects were transported from the adjacent housing room into the procedure room, tails were labeled with a non-toxic permanent marker with the assigned subject ID number, and subjects were left to acclimate undisturbed to the testing environment for a minimum 60 minutes prior to testing. Between subjects, all testing arenas were sanitized with 70% ethanol solution and dried prior to introducing the next subject. Lighting in the testing rooms were consistent with the housing room (∼500 lux) unless where specifically noted. At minimum 5 days post the conclusion of behavioral testing, mice were sent for tissue harvesting.

##### Frailty assessment

Similar to as previously described^15^, subjects were individually evaluated for the absence or presence of 26 aging-related characteristic traits and scored a 0, 0.5, or 1 (based on presence/absence, and severity) for each assessment by a trained observer, blind to genotype/age, and included the following assessments: alopecia; loss of fur color; dermatitis/skin lesions; loss of whiskers; coat condition; piloerection; cataracts; eye discharge/swelling; microphthalmia; nasal discharge; rectal prolapse; vaginal/uterine/penile; diarrhea; vestibular disturbance; vision loss assessed by visual placing upon subject being lowered to a grid; menace reflex; tail stiffening; impaired gait during free walking; tremor; tumors; distended abdomen; kyphosis; body condition; breathing rate/depth; malocclusions; righting reflex. The frailty index score was calculated as the cumulative score of all measures with a maximum score of 26.

##### Core body temperature

Core body temperature was recorded just prior to the conclusion of the frailty assessment via a glycerol-lubricated thermistor rectal probe (Braintree Scientific product# RET 3; measuring 3/4” L .028 dia. .065 tip) inserted ∼2cm into the rectum of a manually restrained mouse for approximately 10 seconds. Temperature was recorded to the nearest 0.1°C (Braintree Scientific product#TH5 Thermalert digital thermometer).

##### Open field activity

Versamax Open Field Arenas (40cm x 40cm x 40cm; Omnitech Electronics, OH USA) were used for this test. Arenas were housed within sound attenuated chambers with lighting in the testing room and arenas consistent with the housing room (∼500 lux). Mice were placed individually into the center of the arena and infrared beams recorded distance traveled (cm), vertical activity, and perimeter/center time. Data were collected in 5- minute time-bins for duration of 60 minutes.

##### Spontaneous alternation

Mice were acclimated to the testing room under ambient lighting conditions (∼ 50 lux). A clear polycarbonate y-maze (in-house fabricated; arm dimensions 33.65cm length, 6cm width, 15cm height) placed on top of an infrared reflecting background (Noldus, The Netherlands), surrounded by a black floor-to ceiling curtain to minimize extramaze visual cues was used for this test. Mice were placed midway of the start arm, facing the center of the y for an 8-minute test period and the sequence of entries into each arm are recorded via a ceiling-mounted infrared camera integrated with behavioral tracking software (Noldus Ethovision XT). Percent spontaneous alternation is calculated as the number of triads (entries into each of the three different arms of the maze in a sequence of three without returning to a previously visited arm) relative to the number of alteration opportunities.

##### Rotarod test for motor coordination

An accelerating Rotarod (Ugo-Basile; model 47600) is used for this test. Lighting in the testing room is consistent with the housing room (∼ 500 lux). The trial began with mice being placed on the rotating rod (4 rpm), which accelerates up to 40 rpm over the course of 300 seconds. Each mouse is subjected to 3 consecutive trials with an ∼1 min inter-trial interval to allow cleaning of the rod between trials. Latency to fall (sec) is measured. Subjects that fall upon initial placement on the rod, before acceleration begins, are scored as 0 sec for that trial.

##### Wheel running activity

Subjects were individually housed into a clean cage with a running wheel (Med-Associates, Vermont, USA) and with food and water ad libitum. The light cycle was identical to the housing room with 12:12 L:D (lights on at 6:00am). Running Wheels were equipped with a wireless transponder that recorded activity on the running wheels (revolutions) in sync with a computer that time stamps events. Mice were left undisturbed throughout the testing period with the exception of daily welfare checks. Data were evaluated for time spent running (min), total distance traveled (meters), and speed (revolutions per min) over the course of three 24-hour periods.

##### Behavioral data analysis

Prior to data analysis and while still blinded, results were adjusted to exclude data only from mice which could not be tested or which data was not available inclusive of any equipment failures, escape episodes, etc. Subjects were not excluded by any mathematical determination. Data was analyzed under coded genotypes (A, B, C, etc.) within sex, as one-way or two-way ANOVA as appropriate versus sex- and age-matched WT control. The blind was revealed at the conclusion of the data analysis for interpretation.

#### Fasted blood glucose collection and measurement

Fasted mice were placed into a fresh cage, free of food but with fresh water, at 6am – the beginning of the light-ON cycle. Mice were fasted for 6 hours, until 12pm, at which time blood glucose levels were analyzed. Prior to mouse restraint, a Contour Next EZ blood glucose monitor (Ascensia, Parsippany, NJ) was calibrated with Contour glucose control solution and Contour Next test strips. While restraining the animal, with a 5.0mm lancet a stab incision was made into and perpendicular to the cheek, located dorsal to the to the cheek skin gland at a distance equal to the height of the eye and caudal distance equal to the length of the eye. One drop of blood, approximately 10μl, was applied to a blood glucose test strip and readings were recorded.

#### Animal anesthesia

Upon arrival at the terminal endpoint for each aged mouse cohort (4, 12, 18 or 24 months), individual animals were weighed prior to intraperitoneal administration of tribromoethanol (1mg/kg). Routine confirmation of deep anesthesia was performed every 5 minutes by toe pinch.

#### Whole Animal Perfusion

First confirming deep anesthetization via toe pinch, an incision is made along the ventral midline to expose the thorax and abdomen, followed by removal of the lateral borders of the diaphragm and ribcage revealed the heart. Prior to perfusion trunk blood and CSF samples were collected. To perfuse the animal, a small cut was placed in the right atrium to relieve pressure from the vascular system before perfusing the animal transcardially with 1XPBS via injection into the left ventricle. Completion of perfusion and clearance of the vascular system was indicated by a blanching of the liver. At this time organs of interest were collected as indicated.

#### Non-fasted blood collection and analysis

Blood was collected by cardiac puncture from non-fasted, anesthetized animals (see Perfusion method) at harvest prior to incision of the right atrium and subsequent perfusion. A 25-gauge EDTA-coated needle, attached to a 1mL syringe, is inserted into the right atrium of the exposed heart and the plunger gently pulled to slowly aspirate approximately 500mL of blood, avoiding entrapping air in the syringe to prevent hemolysis. After removal of the needle from the syringe, the blood was slowly injected into a 1.5mL EDTA coated MAP-K2 blood microtainer (363706, BD, San Jose, CA) on ice. Blood tubes were spun at 4°C and 4,388xg for 15 minutes. Blood serum is then removed and aliquoted equally into three replicate 1.5mL tubes on ice. Tubes were then snap frozen on dry ice and stored long-term at ™80°C. Thawed blood plasma collected from non-fasted mice was then analyzed by Beckman Coulter AU680 chemistry analyzer (Beckman Coulter, Brea, CA) and Siemens Advia 120 (Germany) for levels of non-fasted glucose, total cholesterol, LDL (low-density lipoproteins), HDL (high-density lipoproteins), triglycerides, and NEFA (non-essential fatty acids).

#### Brain harvest

Anesthetized and subsequently perfused animals were decapitated, and heads submerged quickly in cold 1xPBS. The brain was carefully removed from the skull, weighed, and divided midsagitally, into left and right hemispheres, using a brain matrix. The right hemisphere was quickly homogenized on ice and equally aliquoted into three cryotubes for metabolomic, proteomic, and transcriptomic analysis. Cryotubes were immediately snap frozen on dry ice, and stored long-term at ™80°C. The left hemisphere was immediately placed in 5mL 4% PFA at 4°C for no less than 24 hours, but no longer than 30 hours. The left hemisphere was then moved from PFA solution to 10mL 15% sucrose at 4°C for 24 hours, or until it sinks in the sucrose, when it was then transferred to a 30% sucrose for 24 hours at 4°C, or until it sinks in the solution. The left hemisphere was then removed from 30% sucrose solution, snap frozen on a flat mold, cut-side down, floating in 2-methylbutane solution cooled by dry ice. Once frozen the left hemisphere is then placed into a cryotube and stored at ™80°C until used for microtome sectioning and immunohistochemistry analysis.

#### Immunohistochemistry and microscopy imaging

During harvest, whole mouse brains were removed and weighed. Using a brain matrix, left and right hemispheres were separated along the midsagittal plane. The left hemisphere was placed in 5mL of 4% PFA at 4°C overnight, then moved to 10mL of 15% sucrose at 4°C overnight, before finally being incubated in 10mL of 30% sucrose at 4°C overnight or until brain sinks to bottom of the tube. The left hemisphere was then snap frozen and stored at ™80°C until sectioned. Left hemispheres (see preparation in Brain harvest method) were cut via Thermo Scientific HM430 sliding microtome at 25μm thickness. Coronal brain tissue sections were oriented to capture the cortex and hippocampus at approximately Bregma: ™2.75mm and Interaural 1.05mm. Each section was placed into cryoprotectant buffer (37.5% 1xPBS, 31.25% glycerol, 31.25% ethylene glycol) for immediate use or long-term storage at ™20°C. Floating sections were then blocked prior to immunohistochemical staining and mounting. After blocking slides with 10% normal donkey serum or normal goat serum diluted in 1xPBS+0.5%Triton wash buffer all antibodies were washed floating in 1xPBT (1x PBS with 0.5% Triton) wash buffer after blocking for 1 hour at room temperature on shaker in 10% NGS (normal goat serum) or 10% NDS (normal donkey serum) in 1xPBT. Secondary antibodies were incubated in 10% NGS or NDS in 1xPBT for 1 hour at room temperature, followed by washes in 1xPBT before mounting on to slides. Slides were then imaged on a Leica Versa slide scanner, automated fluorescent microscope system (Leica, Allendale, NJ). For further analysis, regions of the cortex and hippocampus were processed using Imaris (Bitplane, Concord, MA) software to quantify cell counts, fluorescence intensity, and surface area ratios. [Iba1 (Wako, 019-19741, 1:300), DAPI (1:1000), GFAP (Origene, AP31806PU-N, 1:1000), NeuN (abcam, ab104225, 1:500), and ThioS (1% in 50% ethanol)]

#### Transcriptomics and Proteomics

RNA-Seq and Tandem Mass Tag Proteomics data were obtained from whole left hemisphere brain samples from mice expressing human *APOEε4* and the *Trem2*R47H* mutation (*APOEε4.Trem2*R47H*; strain name LOAD1) ^8^ and mice expressing humanized Aβ in combination with these genetic risk factors (*APOEε4.Trem2*R47H*.hAβ; strain name LOAD2). Whole-brain left hemispheres were collected at 4, 12, 18, and 24 months of age from both sexes and both diets (standard chow and high-fat, high-sugar diet) where available. Additional C57BL/6J (B6) mice at 18 months were included as a standard aged control. At least six biological replicates were collected for each sample group.

#### RNA-Sequencing data analysis

RNA-Seq data were processed using nf-core/rnaseq pipeline [https://doi.org/10.5281/zenodo.1400710]. Briefly, reads were aligned to the reference mouse genome (version GRCm38.p6) using STAR and gene expression was quantified with RSEM ^42^. To measure human *APOE* gene expression, we created a custom mouse reference genome by concatenating human *APOE* gene sequence (human chromosome 19:44905754-44909393; build GRCh38.p10) into the mouse genome (GRCm38.p6) as a separate chromosome (referred as chromosome 21 in chimeric mouse genome). Subsequently, we added a gene annotation for the human *APOE* gene into the mouse gene annotation file.

#### Differential gene and protein expression analysis

Differentially expressed genes in mouse models were identified using the R Bioconductor package DESeq2 (v1.16.1) ^43^. We used the Benjamini-Hochberg corrected p-values with a significance threshold of 0.05 to identify differentially expressed genes. We performed differential protein expression analysis for each mouse model compared to age and sex-matched control mice using one-way ANOVA followed by post hoc correction using Tukey HSD test to match methods used in analogous human studies ^19^.

#### Functional enrichment analysis

Functional annotations and enrichment analyses were performed using the R Bioconductor package clusterProfiler ^44^, with Gene Ontology term and KEGG pathway enrichment analyses performed using functions enrichGO and enrichKEGG, respectively. The function compareCluster was used to compare enriched functional categories of each gene module. The significance threshold for all enrichment analyses was set to 0.05 using Benjamini-Hochberg adjusted p-values.

#### Weighted gene co-expression network analysis of the mouse transcriptome

Weighted gene co-expression network analysis (WGCNA) was performed to identify modules (clusters) of correlated genes using log TPM normalized expression values. We used the step-by-step construction approach for network construction and module identification ^17^. The default unsigned network type was used, and a soft thresholding power of 5 was chosen to meet the scale-free topology criterion in the pickSoftThreshold function. We set the minimum modules size as 30, and merged modules whose correlation coefficient were greater than 0.75 (mergeCutHeight = 0.25). Each module is summarized by the module eigengene (ME), defined as first principal component of the gene expression profiles of each module. Further we computed the Pearson correlation coefficient of modules to genotype, age, sex, behavioral data, cytokines levels, and cell count data.

#### Enrichment of AD biological domains

Gene functional enrichment analyses are informative but sometimes it is difficult to understand how the enriched terms relate to the biology of AD. Cary et al. developed 19 biological domains that capture the AD-associated endophenotypes and defined them using an exhaustive set of Gene Ontology (GO) terms, with the intent to keep each domain siloed in a biologically coherent fashion. We performed Fisher’s exact tests to identify enrichment of each AD biological domain and resident GO terms in WGCNA module gene sets.

#### Linear regression Analyses

Multiple behavioral data, cytokines levels, NfL, and cell counts were measured for each animal for multiple ages from both sexes. We performed multiple linear regression model to determine the effect of each variant (age, sex, high-fat diet, and genotypes) on these phenotypic data:

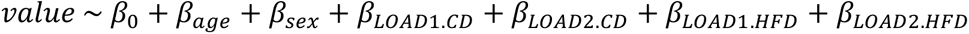

Here the value represents measurements of each quantitative phenotype of interest. CD and HFD refer to chow and high-fat diets, respectively, to quantify the effects of genotypes paired with diet.

#### Proteomics experimental design and data processing

#### Tissue Homogenization and Protein Digestion

Samples were homogenized in 8 M urea lysis buffer (8 M urea, 10 mM Tris, 100 mM NaH2PO4, pH 8.5) with HALT protease and phosphatase inhibitor cocktail (ThermoFisher) using a Bullet Blender (NextAdvance) essentially described ^46^. Each Rino sample tube (NextAdvance) was supplemented with ∼100 μL of stainless-steel beads (0.9 to 2.0 mm blend, NextAdvance) and 300 μL of lysis buffer. Tissues were added immediately after excision and homogenized with bullet blender at 4 °C with 2 full 5 min cycles. The lysates were transferred to new Eppendorf Lobind tubes and sonicated for 3 cycles consisting of 5 s of active sonication at 30% amplitude, followed by 15 s on ice. Samples were then centrifuged for 5 min at 15,000 x g and the supernatant transferred to a new tube. Protein concentration was determined by bicinchoninic acid (BCA) assay (Pierce). For protein digestion, 100 μg of each sample was aliquoted and volumes normalized with additional lysis buffer. Samples were reduced with 5 mM dithiothreitol (DTT) at room temperature for 30 min, followed by 10 mM iodoacetamide (IAA) alkylation in the dark for another 30 min. Lysyl endopeptidase (Wako) at 1:25 (w/w) was added, and digestion allowed to proceed overnight. Samples were then 7-fold diluted with 50 mM ammonium bicarbonate. Trypsin (Promega) was then added at 1:25 (w/w) and digestion proceeded overnight. The peptide solutions were acidified to a final concentration of 1% (vol/vol) formic acid (FA) and 0.1% (vol/vol) trifluoroacetic acid (TFA) and desalted with a 30 mg HLB column (Oasis). Each HLB column was first rinsed with 1 mL of methanol, washed with 1 mL 50% (vol/vol) acetonitrile (ACN), and equilibrated with 2×1 mL 0.1% (vol/vol) TFA. The samples were then loaded onto the column and washed with 2×1 mL 0.1% (vol/vol) TFA. Elution was performed with 2 volumes of 0.5 mL 50% (vol/vol) ACN.

#### Isobaric Tandem Mass Tag (TMT) Peptide Labeling

Each sample (containing 100 μg of peptides) was re-suspended in 100 mM TEAB buffer (100 μL). The TMT labeling reagents (5mg) were equilibrated to room temperature, and anhydrous ACN (256 μL) was added to each reagent channel. Each channel was gently vortexed for 5 min, and then 41 μL from each TMT channel was transferred to the peptide solutions and allowed to incubate for 1 h at room temperature. The reaction was quenched with 5% (vol/vol) hydroxylamine (8 μl) (Pierce). All channels were then combined and dried by SpeedVac (LabConco) to approximately 150 μL and diluted with 1 mL of 0.1% (vol/vol) TFA, then acidified to a final concentration of 1% (vol/vol) FA and 0.1% (vol/vol) TFA. Labeled peptides were desalted with a 200 mg C18 Sep-Pak column (Waters). Each Sep-Pak column was activated with 3 mL of methanol, washed with 3 mL of 50% (vol/vol) ACN, and equilibrated with 2×3 mL of 0.1% TFA. The samples were then loaded and each column was washed with 2×3 mL 0.1% (vol/vol) TFA, followed by 2 mL of 1% (vol/vol) FA. Elution was performed with 2 volumes of 1.5 mL 50% (vol/vol) ACN. The eluates were then dried to completeness using a SpeedVac.

#### High-pH Off-line Fractionation

High pH off line fractionation was conducted essentially essentially described ^46^. Dried samples were re-suspended in high pH loading buffer (0.07% vol/vol NH4OH, 0.045% vol/vol FA, 2% vol/vol ACN) and loaded onto a Water’s BEH 1.7 um 2.1mm by 150mm. An Thermo Vanquish was used to carry out the fractionation. Solvent A consisted of 0.0175% (vol/vol) NH4OH, 0.01125% (vol/vol) FA, and 2% (vol/vol) ACN; solvent B consisted of 0.0175% (vol/vol) NH4OH, 0.01125% (vol/vol) FA, and 90% (vol/vol) ACN. The sample elution was performed over a 25 min gradient with a flow rate of 0.6 mL/min. A total of 192 individual equal volume fractions were collected across the gradient and subsequently pooled by concatenation into 96 fractions and dried to completeness using a SpeedVac.

#### LC-MS/MS Methods

All fractions were resuspended in an equal volume of loading buffer (0.1% FA, 0.03% TFA, 1% ACN) and analyzed by liquid chromatography coupled to tandem mass spectrometry. Peptide eluents were separated on a custom in-house packed CSH 1.7um (15 cm × 150 μM internal diameter (ID) by a Dionex RSLCnano UPLC (ThermoFisher Scientific). Buffer A was water with 0.1% (vol/vol) formic acid, and buffer B was 80% (vol/vol) acetonitrile in water with 0.1% (vol/vol) formic acid. Elution was performed over a 32 min gradient with flow rate at 1000 nL/min. The gradient was from 1% to 99% solvent B. Peptides were monitored on a Orbitrap Eclipse mass spectrometer with a high-field asymmetric waveform ion mobility spectrometry (FAIMS Pro) ion mobility source(ThermoFisher Scientific). Two compensation voltages (CV) were chosen for the FAIMS. For each CV (−45 and ™65) top speed cycle of 1.5 seconds, the full scan (MS1) was performed with an m/z range of 410-1600 at 60,000 resolution at standard settings. The higher energy collision-induced dissociation (HCD) tandem scans were collected at 35% collision energy with an isolation of 0.7 m/z, a resolution of 30,000 with TurboTMT on, an AGC setting of 250% normalized agc target, and a maximum injection time of 54 ms. Dynamic exclusion was set to exclude previously sequenced peaks for 15 seconds within a 10-ppm isolation window.

#### Database Search

Datasets (672 raw files (n=7 batches of 96 high pH fractions)) were searched using FragPipe (version 20.0). The FragPipe pipeline relies on MSFragger (version 3.8; ^47, 48^) for peptide identification and Philosopher (version 5.0.0; ^49^) for FDR filtering and downstream processing. The mouse protein database used contains canonical isoforms from Uniprot/Swissprot as of 02/2023. The workflow used in FragPipe followed default TMT-16 plex parameters, used for both TMT-16 and TMT-18 experimental design. Briefly, precursor mass tolerance was ™20 to 20 ppm, fragment mass tolerance of 20 ppm, mass calibration and parameter optimization were selected, and isotope error was set to ™1/0/1/2/3. Enzyme specificity was set to strict-trypsin and up to two missing trypsin cleavages were allowed. Peptide length was allowed to range from 7 to 50 and peptide mass from either 200 to 5,000 Da. Variable modifications that were allowed in our search included: oxidation on methionine, N-terminal acetylation on protein, and N-terminal acetylation on peptide, with a maximum of 3 variable modifications per peptide. Peptide Spectral Matches were validated using Percolator^50^. The false discovery rate (FDR) threshold was set to 1% and protein and peptide abundances were quantified using Philosopher for downstream analysis.

#### Protein Quantitation and Normalization

The protein abundances are normalized by scaling total protein signal within each channel for each specific case sample to the maximum channel-specific total signal. We then used a tunable median polish approach, TAMPOR, to remove technical batch variance in the proteomic data, as previously described^51^. TAMPOR is utilized to remove intra-batch and inter-batch variance while preserving meaningful biological variance in protein abundance values, normalizing to the median of selected intra-batch samples. This approach is robust to outliers and columns with up to 50% values missing. If a protein had more than 50% samples with missing values, it was removed from the matrix. No imputation of missing values was performed for any cohort. For the current data, TAMPOR leverages the median protein abundance from the pooled Global Internal Standard (GIS) TMT channels as the denominators in both factors to normalize sample-specific protein abundances across the 7 batches of samples.

#### AD-related protein co-expression modules from human postmortem brain tissue

Human protein co-expression modules were obtained from previously published protein expression profiles from human dorsolateral prefrontal frontal cortex ^19^. Briefly, Johnson et al. analyzed more than 500 dorsal prefrontal cortex (DLPFC) tissues from control, asymptomatic AD (AsymAD), and AD brains across multiple centers and cohorts using tandem mass tag mass spectrometry (TMT-MS) based quantitative proteomics and generated a deep TMT AD protein network using WGCNA. This network consists of 44 modules of proteins related to one another by their co-expression across control and disease tissues. Johnson, et al. functionally annotated these modules using Gene Ontology analysis of its constituent proteins and assigned cell types to each module, and measured correlations of each summary module eigenprotein to neuropathological or cognitive traits present in the cohorts ^19^. We obtained case-control LogFC values previously adjusted for sex for each quantified protein and membership in protein co-expression modules from the AD Knowledge Portal (https://www.synapse.org/#!Synapse:syn25453861).

#### Mouse-human correlation analysis

To compare mouse expression changes with those observed in human disease, we computed Pearson correlations between changes in expression (log2 fold change) in human AD cases versus controls and the changes in expression (log2 fold change) in each mouse model contributed by a genotype-diet combination from the linear model (*i.e.* corrected for sex analogous to the human data). Correlations were computed across the set of orthologous proteins in each protein module using cor.test function in R as:

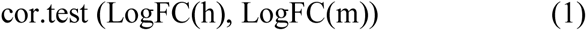

where LogFC(h) is a vector of protein log folds changes in human AD cases compared to controls and LogFC(m) is a vector of protein log fold changes in LOAD mouse models from the linear model.

### Cohort 2 – Indiana University

#### In vivo MR imaging

LOAD2 mice were imaged at the age of 4 months, 12 months and 18 months respectively. For male mice, there were 9 control (CON) and 7 high fat (HF) at 4 months, 9 CON and 10 HF at 12 months, and 7 CON and 10 HF at 18 months respectively. For female mice, there were 10 CON and 9 HF at 4 months, 10 CON and 8 HF, and 8 CON and 4 HF at 18 months respectively.

MR images of the specimens were acquired on a 30-cm bore 9.4 Tesla magnet (Bruker BioSpec 94/30, Billerica, MA, United States) with a maximum gradient strength of 660 mT/m on each axis. A high-sensitivity cryogenic RF surface receive-only coil was used for signal reception (Bruker Cryoprobe). A multi-echo two-dimensional Rapid Acquisition with Relaxation Enhancement (RARE) pulse sequence with 8 echo train length (ETL) was used to acquire T2- weighted images. The scanning parameters were as follows: matrix size = 304 x 256, field-of-view (FOV) = 15.2 x 12.8 mm^2^, in-plane resolution = 100 µm x 100 µm, slice thickness = 250 µm, and repetition time (TR) = 7 s.

#### MRI Data analysis

All Digital Imaging and Communications in Medicine (DICOM) raw data were converted into Neuroimaging Informatics Technology Initiative (NIfTI) format. The mask was created using U-Net^54^. The data were masked and registered to standard T2W template using non-linear diffeomorphic ANTs registration^55^. The template brain labels were warped into subject space following the registration. The mean volume of each label and whole brain volume were calculated, and t-test statistics were conducted to determine statistical significance. The p-value was set at 0.05 level of significance.

#### Perfusion and Preparation of Blood and Tissue Samples

Mice at Indiana University were anesthetized to the surgical plane of anesthesia with tribromoethanol at 4, 12 and 18 months of age. Trunk blood and brain tissue were collected immediately after euthanasia. Trunk blood was centrifuged for 15-20 min at 4°C x 14,500 RPM. Plasma was stored at ™80C.

#### Blood Plasma and Brain homogenate analysis

Blood was collected from non-fasted mice aged 4, 12, 18 months of age. Mice were anesthetized and blood was extracted by left ventricle cardiac puncture with a 25 g EDTA-coated needle before PBS perfusion. Approximately 500 μL of whole blood was transferred to a MAP-K2 EDTA Microcontainer (BD, Franklin Lakes, NJ) on ice and centrifuged at 4°C x 4388 *g* in a pre-chilled ultracentrifuge for 15 minutes. Without disturbing the red blood cell fraction, serum supernatant was pipetted into a chilled cryovial with a with p200 tip and immediately snap-frozen on dry ice for 10 min.

#### Brain homogenization and protein extraction

Hemibrains were homogenized in tissue homogenization buffer containing fresh protease inhibitor cocktail and aliquoted. DEA/Formic Acid extraction was carried out according to Casali et al., 2016. Supernatant was utilized for the cytokine analysis, soluble and insoluble components were used for the Aβ analysis.

#### Cytokine and Aβ Panel Assay

Mouse hemibrain samples were assayed in duplicate using the MSD mouse proinflammatory Panel I, a highly sensitive multiplex enzyme-linked immunosorbent assay (ELISA). The panel quantifies 10 cytokines: interferon γ (IFN-γ), interleukin (IL)-1β, IL-2, IL-4, IL-6, IL-8, IL-10, IL-12p70, IL-13, and tumor necrosis factor α (TNFα) from a single small sample volume (25 μL) using an electrochemiluminescent detection method (MesoScale Discovery, Gaithersburg, MD, USA). The mean intra-assay coefficient for each cytokine was <8.5%, based on cytokine standards. Any value that was below the lowest limit of detection (LLOD) for the cytokine assay was replaced with ½ LLOD of the assay for statistical analysis.

#### Aβ Species Assay

The plasma samples and soluble (DEA) components from hemibrain samples were assayed in duplicate using MSD Aβ Peptide Panel I (K15200E; MSD). Following the manufacturer’s guidelines, the Aβ species (Aβ40 and Aβ42) were quantified from plasma and soluble component of hemibrain lysates using an electrochemiluminescent detection method (MSD). The levels of Aβ species from the assay were used for statistical analysis.

#### Immunohistochemistry

Left brain hemispheres were sectioned at 25 μm on a Thermo Scientific HM430 sliding microtome and blocked with 10% normal donkey or normal goat serum diluted in 1x PBS+0.5% TritonX wash buffer before immunohistochemical staining with antibodies selected to visualize neurons (NeuN), astrocytes (GFAP), microglia (Iba1), and nuclei (DAPI). Images were acquired at 20X using a Leica Aperio Versa (Germany) slide-scanner microscope. Cell counts, fluorescence intensity, and surface area ratios in the regions of the cortex and hippocampus were calculated using Imaris (Chicago, IL, USA).

### Cohort 3 – Indiana University

To assess neurovascular coupling, mice were non-invasively imaged via PET/CT (n=12 mice/sex/genotype/age). Regional blood flow was be measured via ^64^Cu-pyruvaldehyde-bis(N4- methylthiosemicarbazone) (^64^Cu-PTSM) ^56^, which has a very high first pass (>75-90%) extraction^57^, and glutathione reductase redox trapping of copper^57^, was administered via tail vein in awake subjects, and allowed 2 min uptake period. To measure regional glycolytic metabolism, 2-^18^F-2-deoxyglucose (^18^F-FDG) was administered via inter-peritoneal injection in awake subjects and mice were given 30-45 min uptake period prior to imaging per our previous work ^8^. Post uptake, mice were induced with 5% isoflurane (95% medical oxygen) and maintained with 1-2% isoflurane at 37°C. PET/CT imaging were performed with a Molecubes β-X-CUBE system (Molecubes NV), where calibrated listmode PET images were reconstructed into a single-static image previously described^8^. Helical CT images were also acquired for anatomical reference, and attenuation maps needed to correct PET images per our previous work^8^. PET and CT images were co-registered, and mapped to stereotactic mouse brain coordinates of Paxinos-Franklin^58^. To permit dose and scanner and brain uptake normalization, Standardized Uptake Value Ratios (SUVR) relative to the cerebellum were computed for PET for each subject, genotype, and age as follows:

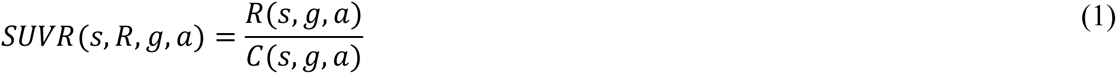

where, *ss*, *gg*, *vv*, *VV*, and *CC* are the subject, genotype, age, region/volume of interest, cerebellum region/volume of interest. The SUVR values were then converted to z-score as follows:

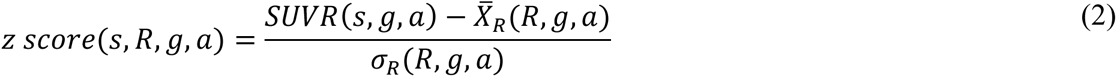

where, *s*, *g*, *v*, *V*, *X_R_* and *σ*_*R*_ are the subject, genotype, age, mean of the reference population in SUVR, standard deviation of the reference population, based on the specified analytical strategies (effects of aging, humanized genes, and AD-risk alleles). Data are then plotted on a Cartesian coordinate system, where [0,0] is no change in tissue perfusion (x-axis) or metabolism (y-axis) from the reference population.

#### Cryotomy

After PET imaging, animals were euthanized and brains were removed, bisected, and snap-frozen on dry ice before embedding in OCT in cryo-molds. Brains were sectioned at 20 μm with a Leica 1850 or CM1860 Cryotome, with 6 per slide per bregma region of interest. Bregmas: 0.38, ™1.94, ™3.8, ™5.88. ROIs: Corpus Callosum, Striatum, Cing Ctx, Motor Ctx, Somatosensory Ctx, DI Ctx, Med Sep, Hypothalamus, Hippocampus, Retrosplenial Ctx, Auditory Ctx, Entrohinal Ctx, Thalamus, TEA, Visual Ctx, Cerebellum.

#### Autoradiography

Standards for calibration were prepared using two-fold serial dilutions of PET tracer in 3.2% low melt agarose, 4% sucrose, 0.2% gelatin. Aliquots (∼30 μL) were dotted into cryomolds and allowed to firm up. OCT was added and standards were frozen and cut at the same thickness as the brain sections. An additional ∼20 μL aliquot was pipetted into a 7 mL scintillation tube filled with scintillation fluid and counted on a Beckman LSC 6500 for 15 to 30 seconds using the wide window. These data were used to calculate the standard curve for each animal and were converted to Bq/ml using a standard calculation worksheet. Slides with brain sections were placed on cardboard along with standards and exposed overnight to phosphorimager before imaging on the GE Typhoon FLA7000IP.

### Cohort 4 – University of Pittsburgh

#### Breeding and Husbandry

After weaning, all mice were housed by sex, 2-5 mice per cage with P.J. Murphy Coarse Certified Aspen Sani-Chip® bedding and initially provided LabDiet® 5P76 (6% fat; control diet, CD) *ad libitum* until 2 months of age. At 2 months of age, mice assigned to the HFD group were given *ad libitum* access to a 45% kcal fat rodent diet (Research Diets, D12451i). All cages included enrichment consisting of nestlets (Ancare, NES3600) and red PET plastic domes (Braintree Scientific). The mice were kept on a 12-hour light/dark schedule with the lights on from 7:00 a.m. to 7:00 p.m. daily. The room was maintained at 72-74°F and 30-70% humidity. Chlorinated water (2-4ppm; pH 4-5) was provided *ad libitum* via a lixit system. The animals were ear-punched for identification purposes and subsequently microchipped at the tail base using a p-chip system (PharmaSeq).

#### Plasma biomarkers

#### Cytokines

Subjects were evaluated monthly for plasma cytokines from 2-13 months of age and then at 15 and 18 months of age (Figure 8A). Plasma was also collected and analyzed at 8.5 and 10.5 months, each after 2-week bouts of food restriction as described in the following sections. For longitudinal analysis of plasma cytokines, 50µL blood was collected via the tail tip in non-anesthetized mice into heparinized capillary tubes and then transferred to chilled 1.5mL microtubes and centrifuged @4C for 10 min x 14500 rpm. Plasma was stored as 25µL aliquots at ™80C until analysis. MesoScale Discovery multiplex ELISA kits were run according to the manufacturer’s protocol for the mouse pro-inflammatory cytokine panel (Kit# K15048D).

#### Aβ40 and Aβ42

Subjects were evaluated at 2, 6, 9, 12, 15, and 18 months for plasma Aβ40 and 42 (Figure 8A). After a heparinized blood sample was collected for analysis of plasma cytokines, 40-50µL blood was collected via the same tail tip into EDTA coated capillary tubes and then transferred to chilled 1.5mL microtubes and centrifuged @4C for 10 min x 14500 rpm. Plasma was stored as 20µL aliquots at ™80C until analysis. MesoScale Discovery (MSD) multiplex ELISA kits were run according to the manufacturer’s protocol for the 6E10 Aβ peptide panel (Kit# K15200E).

#### Statistical Analysis of Plasma Biomarkers

MSD ELISA results were analyzed in Discovery Workbench Software provided by MSD. All samples from a single plasma collection timepoint were run together on the appropriate MSD plate with calibration standards. The calibration standards from each plate and the accuracy and precision between and within runs were evaluated based on the FDA’s M10 Bioanalytical Method Validation guidelines for ligand binding assays. Calibration standards whose back-calculated concentrations exceeded ±20% (or ±25% at the ULOQ and LLOQ) of the expected concentration (% recovery) were excluded. At least 75% of all calibration standards and at least 6 concentrations per assay were required to meet criteria for each plate to be included in statistical analysis. Across all plates run for each assay, the % recovery for each included calibration standard ranged from 95-105%.

After completion of within and between plate quality control, plasma biomarker concentrations for 8 cytokines (TNF-α, IL-6, IL-10, IL-1β, IFN-γ, IL-5, KC/GRO, and IL-2), Aβ40, and Aβ42 were compared using two-way ANOVA with Tukey’s multiple comparisons test and a 95% confidence interval.

#### Food Restriction

Beginning at 8 months of age, mice were individually housed and restricted to 80-85% of free-feeding body weight (Figure 8A). Mice were weighed daily and provided a ration of the respective CD or HFD diets that maintained them at 80-85% restriction.

#### Touchscreen cognitive testing

Male and female LOAD2 mice exposed to ad libitum high fat diet (HFD) from 2 months of age (Figure 8A) and LOAD2 or C57BL/6J (WT) mice exposed to HFD from 6+ months of age (Supplemental Figure 7A) and age- and sex-matched LOAD2 and C57BL/6J WT controls maintained on normal control diet (CD) chow were enrolled for cognitive testing beginning from after 12 months of age. Prior to enrollment blood was collected (non-anesthetized) up to twice per month via the tail collection method as described above; otherwise, subjects were behaviorally naïve prior to touchscreen task acquisition. Mice were trained and tested in daily sessions (typically 5-6 days per week) using the Bussey-Saksida mouse touchscreen chambers and ABET II software pre-programmed protocols for the Location Discrimination task (Lafayette Instrument Company, Lafayette, IN, USA) similar to the methods previously described for the location discrimination task^11^. Trained technicians were blinded to genotype throughout testing though blinding for diet was not possible due to visual differences in HFD v CD. Genotype, sex, and diet treatment were randomized and counterbalanced across multiple chambers and sessions though the same subject was always assigned to the same chamber throughout training and testing. The touchscreen chambers were enclosed in sound-attenuated and ventilated chambers and a black Perspex mask consisting of two rows, each of 6 adjacent square response windows placed directly in front of the touchscreen was used to isolate only the areas on the touchscreen that are relevant to the task stimuli presented and minimize non-targeted touch responses to the screen. During the pre-training phase, subjects were food restricted to 80- 85% of free feeding body weight and acclimated to the reinforcer used for the training and testing in their homecages (10% sucrose solution formulated in drinking water). Once subjects were within the requisite restricted weight range, habituation and training using a step wise criterion-based approach was initiated similar to as previously described^11^). During the initial acquisition phase of the task, subjects were trained to associate nosepoke touches to an illuminated square on the screen with the presentation of a reward (20µL of 10% sucrose solution). Touch responses to blank squares had no programmed consequences. This ‘must touch’ phase was subsequently followed by a ‘punish incorrect’ phase in which touch responses to illuminated stimuli were rewarded, while touches to blank squares resulted in the illumination of the houselight with no reward delivered. Accuracy as an indicator of learning was analyzed during the punish incorrect phase of the training by two-way repeated measures ANOVA (treatment group x time). Subjects that failed to meet criterion during each step were not advanced for further cognitive testing. More specifically, subjects that failed to meet task acquisition criteria during the punish incorrect phase were not assessed for pattern separation in the location discrimination (LD) task. For subjects that advanced to the LD task, during LD acquisition trials only the bottom row of 6 adjacent squares was used and annotated for description purposes as squares 1-6 from left to right (see Oomen et al 2013). During the LD training phase, only squares 2 and 5 were used that represent an intermediate location separation for left and right positions, respectively on the screen. During the Intermediate sessions, both left and right squares were illuminated with one designated as correct and the other as incorrect. A nosepoke response made to the correct location is rewarded while incorrect responses result in a 5 second timeout period paired with the houselight turning on. After a 10 second inter-trial interval (ITI), both stimuli in the identical locations is presented again. Once the subject makes 7 correct out of 8 consecutive responses, then the rule is reversed and the subject must determine through trial and error the correct rewarded location. For each consecutive session, the location (left or right) assigned as correct at the start of the session was reversed to avoid mediating strategies. A priori criterion to advance to further pattern separation assessments is 2 consecutive days with ≥3 serial reversals within a session (max trials allowed, max session time limit = 60 min). Upon meeting criteria during the intermediate phase of the test, subjects are evaluated in 2 consecutive sessions with the locations of the two stimuli with the furthest separation = “easy” difficulty (positions 1 and 6) followed by 2 consecutive sessions with the closest separation of the two stimuli = hard difficulty (positions 3 and 4). The primary measures of performance are the # trials to criterion.

### Cohort 5 – University of Pittsburgh

#### Breeding and Husbandry

After weaning, all mice were housed by sex, 2-4 mice per cage with P.J. Murphy Coarse Certified Aspen Sani-Chip® bedding and initially provided LabDiet® 5P76 (6% fat; control diet, CD) *ad libitum* until 6-12 months of age. At 6-12 months of age, mice assigned to the HFD group were given *ad libitum* access to a 45% kcal fat rodent diet (Research Diets, D12451i). All cages included enrichment in the form of nestlets (Ancare, NES3600) and red PET plastic domes (Braintree Scientific). The mice were kept on a 12-hour light/dark schedule with the lights on from 7:00 a.m. to 7:00 p.m. daily. The room was maintained at 72-74°F and 40-60% humidity. Chlorinated water (2-3ppm; pH 4-5) was provided *ad libitum* via a lixit system. The animals were ear-punched for identification purposes and subsequently microchipped at the tail base using a p-chip system (PharmaSeq). Prior to beginning touchscreen training at 11-17 months of age (Supplementary Fig 7A), mice were individually housed and restricted to 80-85% of free-feeding body weight. Mice were weighed daily and provided a ration of the respective CD or HFD diets that maintained them at 80-85% restriction.

#### Plasma biomarkers

#### Cytokines

Subjects were evaluated monthly for plasma cytokines from 6-12 until 11-17 months of age (Supplemental Figure 7A). Collection, storage, and analysis of plasma for proinflammatory cytokines followed the methods described above for Cohort 4.

#### Aβ40 and Aβ42

Subjects were evaluated at a single timepoint, 11-17 months, for plasma Aβ40 and 42 (Supplemental Figure 7A). Collection, storage, and analysis of plasma for Aβ species followed the methods described above for Cohort 4.

## Notes

https://doi.org/10.7303/syn53128146

### Competing Interest Statement

The authors have declared no competing interest.

